# Genome-wide diversity across 130 years reveals drastic post-bottleneck genetic erosion in a recovering population of golden eagles *(Aquila chrysaetos)*

**DOI:** 10.64898/2026.11.06.764905

**Authors:** Ekaterina Karabanina, Suvi Ponnikas, Tiina M. Mattila, Alina Niskanen, Laura Kvist

## Abstract

Many wild populations have experienced severe human-induced declines, but the genetic consequences of such bottlenecks are often poorly understood. In Finland, as well as in many other countries, golden eagles were heavily persecuted during the 19^th^-20^th^ centuries, disappearing from southern regions and reaching their lowest population size in the 1950–1960s. Although conservation efforts facilitated demographic recovery, the long-term genomic consequences of the bottleneck remained unknown. Here, we used whole-genome sequencing data spanning 136 years to evaluate patterns of genetic diversity in 26 Finnish golden eagles before, during, and after the mid-20^th^ century bottleneck. We found substantial loss of genetic variation, including an 82% reduction in genome-wide heterozygosity, a 42% decline in nucleotide diversity, a 67% reduction in Watterson’s theta, and a substantial loss of variable sites. Analyses of the site frequency spectrum and Tajima’s D confirmed historical population contraction, and runs of homozygosity analysis indicated elevated post-bottleneck inbreeding. These results indicate strong genetic erosion in Finnish golden eagles despite the census size recovery. Therefore, incorporating genomic indicators in conservation monitoring and performing regional assessments are necessary to guide management and preserve biodiversity in line with the Global Biodiversity Framework’s goal of maintaining genetic variation.

## Introduction

Human activities have strongly influenced natural environments, causing severe population declines (i.e., population bottlenecks), fragmentation, and extinctions in many species (Ceballos et al., 2015; Dirzo et al., 2014; Young et al., 2016). Beyond their immediate demographic effects, population bottlenecks can leave long-lasting genetic consequences that compromise the adaptive potential and long-term viability of species via elevated genetic drift, increased inbreeding, accumulation of deleterious mutations, and overall higher genetic load (Díez-del-Molino et al., 2018). Because of its role in evolutionary potential (Frankham, 2022), genetic diversity is considered a key component of biodiversity (United Nations Convention on Biological Diversity, 2022). However, understanding whether populations maintain genetic diversity requires quantitative assessment (Hoban et al., 2020). In this context, genetic (or genomic) erosion describes the loss of genetic diversity and evolutionary potential of populations, as well as the loss of structures and processes that maintain genetic resilience, such as connectivity among populations (Hoban et al., 2020). Genetic erosion can result from natural or anthropogenic processes, including habitat fragmentation, population bottlenecks, diseases, hybridization, and inbreeding (Hoban et al., 2020). However, ancient bottlenecks and prolonged small population sizes can mask the genetic effects of recent declines (Díez-del-Molino et al., 2018; Dussex et al., 2015; Hoffman & Blouin, 2004). Historical samples offer a way to address this challenge, allowing researchers to distinguish long-term low diversity from more recent genomic erosion, and museum collections are an invaluable source of such samples (Nakahama, 2021).

The golden eagle (*Aquila chrysaetos*) is a large raptor with a wide Holarctic distribution (Watson, 2010). The species was widespread throughout Finland (except the Åland Islands) until the 19^th^ century, but its distribution and abundance declined sharply due to intensive human persecution (Figure 1) (Ollila, 2018). The available but limited historical records show that already in 1741, bounties were paid for killing “pest birds” (birds of prey), and for collecting their eggs (Vuorisalo & Laihonen, 2000). This persecution was later formalized, and in 1868 large carnivores and raptors were classified as harmful species under Finnish hunting legislation, leading to sustained killing of golden eagles until the early 20^th^ century (Erkamo, 1990; Vuorisalo & Laihonen, 2000). These actions resulted in a population collapse, with the most credible estimate of 250–300 breeding pairs remaining in the 1950s and 1960s – the lowest population size for two centuries (Koskimies & Ollila, 2024). The true number of golden eagles during that time remains uncertain due to incomplete knowledge of territories and lack of systematic monitoring. However, it is known that during this period, the eagles disappeared from southern Finland (Figure 1) (Ollila & Koskimies, 2007) and that breeding success reduced by 37%, estimated as the number of young per occupied territory during 1976–1980, representing the lowest level recorded in the subsequent thirty years (Koskimies & Ollila, 2024). Systematic monitoring began only in the 1970s, after the population collapse, and therefore the pre-bottleneck population size remains unknown.

**Figure 1.**
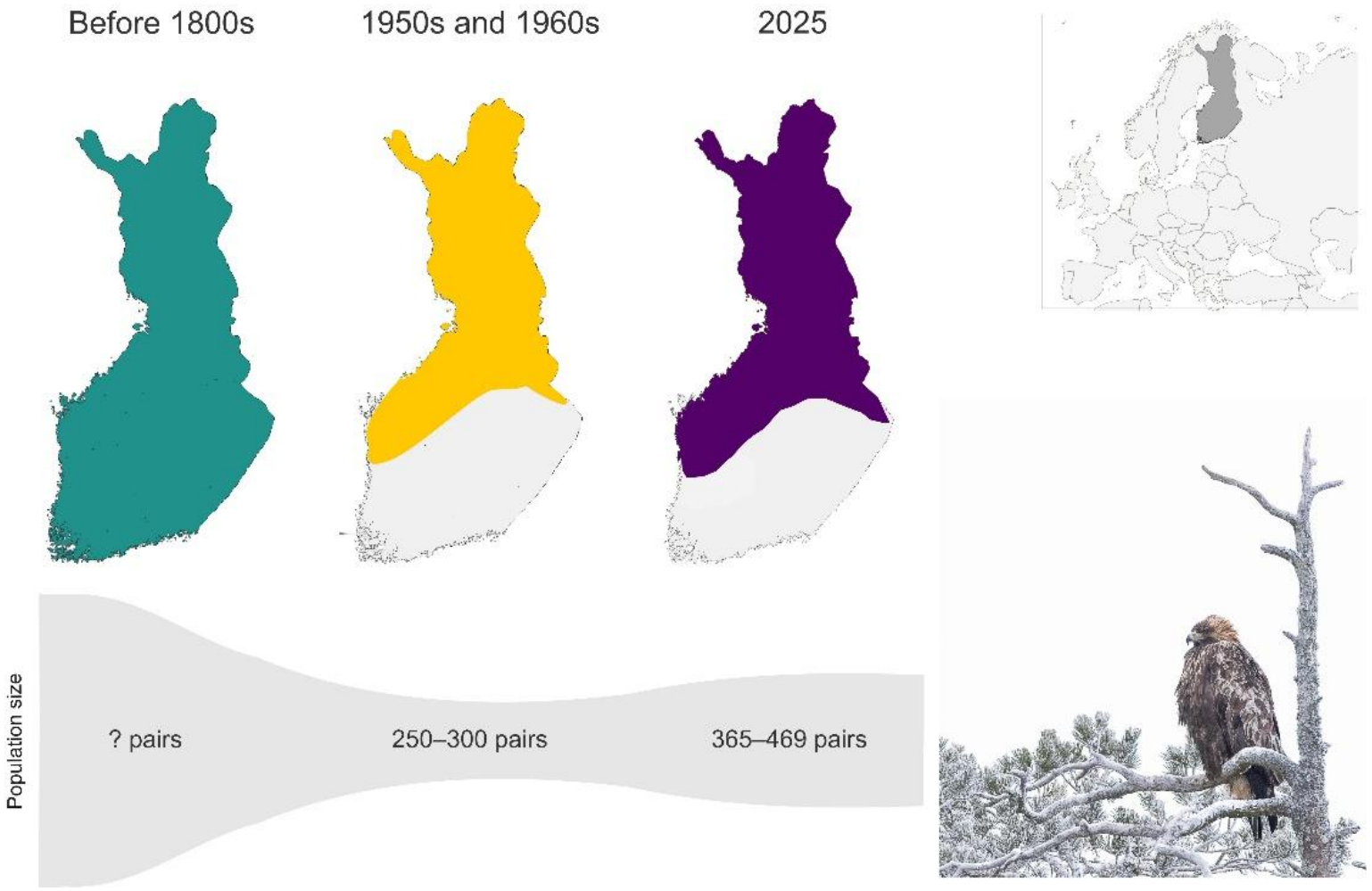
Golden eagle distributions and breeding population sizes in Finland during the three temporal periods in this study: pre-bottleneck (green; before the 19th century), bottleneck (yellow; 1950s–1960s), post-bottleneck or modern (purple; 2025). The top right panel shows the location of Finland on a map of Europe. Photo of the golden eagle by Olli-Pekka Karlin.

In response to population decline, legal protection was introduced gradually: first in southern Finland in 1926, and later in the reindeer herding area in northern Finland in 1962 (Below, 2000). However, permits to kill golden eagles were still issued in the reindeer herding area, and full protection was granted nationwide only in 1969 (Below, 2000). Illegal killing and nest destruction nevertheless continued into the early 2000s (Koskimies & Ollila, 2024). In addition to protection (golden eagles, their nest sites, and eggs; Nature Conservation Act 9/23), targeted conservation measures, such as a compensation scheme for semi-domestic reindeer depredation (Kvist et al., 2026), the construction of artificial nests, and supplementary winter feeding have facilitated the demographic recovery of the golden eagle. Current monitoring estimated 365–469 breeding pairs in Finland in 2025, with over 80% of territories located in the northern partc of the country (Figure 1) (Sundvall, 2025). The species is listed as Vulnerable in The Red List of Finnish species (Hyvärinen et al., 2019).

Despite the demographic recovery and the absence of notable changes in genetic diversity over the last 15 years (Kylmänen et al., 2023), a broader study across Eurasia showed that golden eagles experienced reduced genetic diversity following the 20^th^-century population bottlenecks across the Palearctic range (Karabanina et al., 2024). However, that analysis relied on a limited number of genetic markers, which may constrain the resolution of genome-wide patterns of genetic change. Furthermore, the results were based on a pooled dataset of individuals from across Eurasia. Given that genetic diversity of golden eagles varies across the Palearctic range (Karabanina et al., 2024; Nebel et al., 2015, 2019), regional patterns of genetic change remain unknown. Finally, although the North European population was found to have lower genetic diversity compared to the rest of the species’ Palearctic distribution (Karabanina et al., 2024), it is unclear whether it resulted from regional historical bottlenecks or was a consequence of large-scale demographic processes, such as slower recolonization (Eckert et al., 2008).

In this study, we explored the genomic consequences of the mid-20^th^ century population bottleneck of golden eagles in Finland. We compared low-coverage genome-wide data from three temporal groups: before the bottleneck (1886– 1914), during the bottleneck (1955–1969), and the modern population (2012– 2021). Specifically, we assessed (1) genome-wide genetic diversity, (2) genetic differentiation among time periods, (3) inbreeding and runs of homozygosity, and (4) changes in demographic history. The results of this study quantified the extent of genetic erosion in golden eagles in Finland following persecution and highlighted the importance of incorporating genetic indicators into conservation assessment of demographically recovering populations.

## Materials and methods

### Sampling, DNA extraction, and whole-genome re-sequencing

Tissue samples (toe pads, skin, and muscle tissues) were collected from 30 golden eagle specimens at the Finnish Museum of Natural History and the Oulu Zoological Museum, representing three temporal periods: pre-bottleneck (years 1886–1914; hereafter PRE, N = 10), bottleneck (1955–1969; BOT, N = 10), and post-bottleneck or modern (2012–2021; POST, N = 10) (Table S1). DNA from modern samples was extracted using the E.Z.N.A. Tissue DNA Extraction Kit (Omega Bio-tek, Inc., Norcross, GA, USA), following the manufacturer’s protocol. DNA from historical samples (PRE and BOT) was extracted using a custom phenol-chloroform protocol in a dedicated clean laboratory to minimize contamination (Appendix 1).

DNA concentration was quantified with PicoGreen (Thermo Fisher Scientific, Waltham, MA, USA), and purity was assessed using a NanoDrop spectrophotometer (Thermo Fisher Scientific, Waltham, MA, USA) based on the 260/280 ratio. Of each DNA extract, 30 μl were sent to Novogene for library preparation (350 bp insert size) and Illumina (paired-end, 150 bp) whole-genome sequencing. The target yield was 12 Gb of raw data per sample, corresponding to approximately 10× coverage given the golden eagle genome size of 1.23 Gb (Mead et al., 2021). Three PRE samples failed during library preparation and were excluded from further processing and analyses.

### Read quality control and mapping

A block scheme and scripts of bioinformatic analysis steps are presented in Appendix 2. All subsequent statistical tests and plotting were performed in R v4.5.1. (R Core Team, 2021) using RStudio 2025.05.0. (Posit Software, PBC).

Initial quality control and read mapping of the 27 successfully sequenced samples were performed by Novogene. Raw reads were processed with fastp v0.20.0 (Chen et al., 2018). PolyG tails were removed (-g), and reads were excluded if the base quality was below 5 (-q 5), if they contained more than 15 undetermined bases (-n 15), or if more than half of their bases were unqualified (-u 50). Reads shorter than 150 bp were discarded (-l 150). One mismatch in paired-end reads was allowed (-overlap_diff_limit 1), and overlapping regions were identified if at least 90% of bases matched (-overlap_diff_percent_limit 10). Adapter sequences were automatically detected and removed by fastp. Filtered reads were aligned to the golden eagle reference genome (NCBI GenBank accession number GCA_900496995.4) using the BWA-MEM algorithm v0.7.8 (H. Li, 2013) and the alignments were sorted with samtools v1.3.1 (Danecek et al., 2021). BAM files of the same individual that were run twice were merged with Picard v1.111 using the MergeSamFiles option (Broad Institute). Duplicate reads were removed with samtools rmdup, and final alignments were indexed with samtools.

After mapping, sequencing depth and coverage statistics were obtained using samtools depth and a custom Bash script. Realignment around indels was performed with GATK v3.8.1 (Van der Auwera & O’Connor, 2020). To account for post-mortem DNA damage, base quality score recalibration was done using MapDamage2 (Jónsson et al., 2013). Only autosomal alignments were retained, and soft-clipped reads were excluded. Sequencing depth and breadth were recalculated on the final BAM files. One sample from the BOT group (ACH76) that showed low sequencing coverage (35.5% breadth; 1.37× depth) was excluded from downstream analyses.

Substitution error rates were estimated for the final dataset of 26 samples using the -doError function in ANGSD v0.940 (Korneliussen et al., 2014) based on genotype likelihoods (GL) and applying the same quality and mapping settings as those used for GL estimation (see section “Genotype likelihoods and linkage disequilibrium pruning”), except that a minimum requirement of 10,000 callable sites per individual (-minSites) was applied. SNP calling parameters such as -snp_pval were not used, because error rate estimation was performed directly on allele counts (-doMajorMinor 2).

### Sequencing depth standardization and robustness evaluation

Genome-wide sequencing depth varied among samples (2.08×–9.73×; Table S1) and showed a correlation with some population genetic estimators (e.g., heterozygosity; Spearman’s *ρ* = -0.76, p = 1.0 × 10^-5^). To standardize coverage across samples, all BAM files were downsampled to 2× using the samtools view -S option. For downsampling, a fraction of random reads was retained to achieve an average sample sequencing depth of 2×. Analyses presented below are based on this downsampled dataset.

To evaluate the robustness of our results to data filtering and sequencing depth, we compared the main dataset (downsampled to 2× with transitions) with two supplementary datasets: (1) downsampled to 2× excluding transitions and (2) the full-depth dataset including transitions (Table S2). Obtained results on temporal patterns were consistent across all datasets (Appendix 3, Table S3).

### Genotype likelihoods and linkage disequilibrium pruning

Genotype likelihoods (GLs) were estimated for the total dataset (N = 26) in ANGSD using the samtools model (-GL 1). Major and minor alleles were inferred from GLs (-doMajorMinor 1), assuming the major allele corresponds to the reference genome, while the minor allele was inferred by summing over the probabilities of the three possible alternatives (-doMaf 2). A minor allele frequency threshold (-minMaf) of 0.02 was applied to exclude singletons. A minimum mapping quality (-minMapQ) was set to 30, and a minimum base quality (-minQ) to 20. Base alignment quality computation and mapping quality adjustment were enabled (baq -1, -C 50). Only uniquely mapped, properly paired, and high-quality (flags below 255) reads were retained (-uniqueOnly 1, -only_proper_pairs 1, -remove_bads). A site was considered variable if it was observed in at least five individuals (-minInd 5), corresponding to approximately 70% of the smallest size group (PRE, N = 7). This threshold was selected to retain rare alleles in PRE samples that might have been missed by using a threshold based on a fraction of the total dataset. Minimum and maximum read depths per individual (-setMinDepthInd, -setMaxDepthInd) were set to 1 and 8 (4 × average depth to exclude variable sites within repetitive regions), respectively. Only biallelic SNPs (-skipTriallelic 1) with a p-value threshold (-snp_pval) of 1 × 10^-6^ were called. Transitions were either kept (for runs of homozygosity analyses) or excluded (-rmTrans 1; for PCA and Admixture analyses), and output likelihoods were saved in Beagle GL format (-doGlf 2).

Linkage disequilibrium (LD) was estimated for transversion SNPs using ngsLD (Fox et al., 2019) for all SNP pairs within 500 kb. An LD-decay curve was generated from a random 10% of the data, binned into 150 intervals (Figure S1). Based on the decay pattern, SNPs within 150 kb showing R^2^ > 0.12 were pruned using prune_graph (Vieira, 2025). The resulting set of LD-pruned transversion SNPs was used for PCA and Admixture analysis.

### Site frequency spectrum

The genome-wide site frequency spectra (SFS) produced with ANGSD were used as a basis for estimating genetic diversity and population differentiation (see below). A folded SFS, also known as the minor allele frequency SFS, was chosen because the ancestral allele could not be inferred. To produce SFS, ANGSD first calculated site allele frequency (SAF) likelihoods, and the realSFS subprogram optimized these likelihoods by expectation-maximization to produce folded spectra. SAF likelihoods were calculated from genotype likelihoods (-doSaf 1) using the samtools model (-GL 1) and applying the same quality filters across all analyses (-remove_bads 1, -uniqueOnly 1, -only_proper_pairs 1, -baq 1, -C 50, -minMapQ 30, -minQ 20, -doCounts 1, -setMinDepthInd 1, -setMaxDepthInd 8). Triallelic sites were automatically excluded, and no minor-allele-frequency filters were applied to avoid distorting the SFS (Matz, 2018). To assess the robustness of SFS-based results, analyses were repeated excluding transitions by supplementing the options -doMajorMinor 1 and -rmTrans 1 in the ANGSD commands for SAF estimation.

Four folded SFS types were generated for downstream analyses: (1) per-individual SFS, (2) one-dimensional (1D) SFS per group using original sample sizes, (3) 1D SFS per group downsampled to equal sample size (N = 7), and (4) pairwise 2D SFS. Downsampling the sample sizes was applied to ensure comparability across groups in analyses that directly used the SFS shape, because the SFS shape is affected by the number of gene copies. Groups were therefore reduced to the smallest sample size (N = 7) by removing individuals with the lowest breadth of coverage (Table S1). For the downsampled 1D SFS, the minimum-individual requirement for producing SAF (-minInd) was set to 5 for all groups, and for other types of SFS, it was set to 5 for PRE, 6 for BOT, and 7 for POST. This downsampled 1D SFS was visualized in R to examine the distribution of minor allele frequencies and to check for signals of population bottlenecks.

### Genetic diversity

#### Heterozygosity

Genome-wide heterozygosity was estimated from per-individual folded SFS, each generated with 100 bootstrap replicates. Individual heterozygosity was calculated as the proportion of polymorphic sites among all sites in the SFS output. Group-level heterozygosity and standard errors were calculated from these values. Data normality was assessed using the Shapiro-Wilk test, and, as normality assumptions were not met, group comparisons were performed using non-parametric tests: Kruskal–Wallis rank-sum test for multiple groups and Mann-Whitney U test (Wilcoxon rank-sum) for pairwise comparisons. The relative loss of heterozygosity between PRE and POST groups was quantified as ΔH following Díez-del-Molino et al. (2018) :

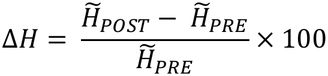

where *H̃* is median heterozygosity. Medians were used instead of averages due to non-normal data distributions.

#### Nucleotide diversity, Watterson’s theta, Tajima’s D, and proportion of variable sites

Nucleotide diversity (π), Watterson’s theta (θ_W_), and Tajima’s D were estimated per group in ANGSD using 1D folded SFS on full sample sizes of each group. SAF likelihoods were transformed into per-site thetas (saf2theta command), and chromosome-level summaries were obtained with the thetaStat subprogram using the empirical Bayes method. For each chromosome, ANGSD returned the sum of θ across callable sites. Genome-wide π and θ_W_ were calculated by weighting the chromosome’s estimates by the number of callable sites as:

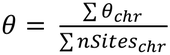

where *θ* is either π or θ_W_, *θ_chr_* is π or θ_W_ per chromosome, and *nSites* is the number of callable sites in each chromosome. The relative loss of nucleotide diversity (Δπ) and Watterson’s theta (Δθ_W_) between the PRE and POST groups were calculated using the same approach as for ΔH. Due to the non-normal data distribution, across-group statistical comparisons were performed using the Friedman test with chromosomes set as dependent variables. To visualize the distribution of Tajima’s D across the genome using Manhattan plots, analyses were repeated using non-overlapping 20 kbp windows (thetaStat do_stat command supplied with -win 20000 -step 20000).

Unlike the estimates above, the proportion of variable sites was calculated from the downsampled 1D SFS for each group by dividing the number of segregating sites by the total number of sites.

### Genetic structure, admixture, and differentiation

Genetic differentiation, admixture, and clustering of individuals from the three temporal groups were assessed with Principal component analysis (PCA), admixture inference, F_ST_, and population branch statistics (PBS).

PCA was performed with pcangsd (Meisner & Albrechtsen, 2018) on the LD-pruned transversion SNPs. A minor allele frequency (--maf) threshold was set to 0.02 to exclude singletons, and the algorithm was iterated 5000 times to achieve convergence. The number of principal components was determined by pcangsd using the minimum average partial (MAP) test.

Admixture analysis was performed with the ngsAdmix program (Skotte et al., 2013) using the same SNP set as in PCA. Runs were performed for 1 to 5 clusters (K), each repeated 100 times, with a minor allele frequency (-minMaf) filter of 0.02 and including SNPs present in at least five individuals (-minInd 5). The most likely number of clusters was inferred from the log-likelihood (LnP(K)) and Evanno’s ΔK (Evanno et al., 2005) statistics, calculated with a custom script in R. Admixture proportions were visualized using the pophelper package (Francis, 2017) in R.

Pairwise F_ST_ and PBS were estimated from 2D folded SFS generated in ANGSD. While F_ST_ quantifies genetic differentiation between pairs of groups, PBS allows identification of the specific group contributing most to the observed differentiation. Thus, for each population pair, joint 2D-SFS priors were inferred in realSFS, which then calculated F_ST_ and PBS in non-overlapping 20 kbp windows (-win 20000 -step 20000). Genome-wide weighted F_ST_ for each population pair was calculated by weighing the per-window F_ST_ estimates by the number of callable sites:

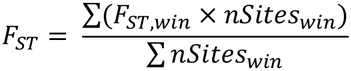

where F_ST_ is the genome-wide F_ST_ for a population pair, F_ST,win_ is the F_ST_ per window, and nSites_win_ is the number of callable sites in a window. Weighted PBS were calculated in the same manner.

PBS results were visualized as Manhattan plots to identify regions of high genetic differentiation. To assess the robustness of the F_ST_ and PBS analyses, 20 permutations were performed by randomly reassigning individuals to three groups while maintaining original sample sizes (PRE = 7, BOT = 9, POST = 10) and re-running the ANGSD analyses (doSaf, realSFS, fstIndex, and fst stats2) for each permutation. The observed weighted F_ST_ and PBS values were compared against the distributions obtained from the permuted datasets.

### Inbreeding and runs of homozygosity

Runs of homozygosity (ROH) were estimated as a measure of population inbreeding using the RZooRoh package v.0.4.1 (Bertrand et al., 2019; Druet & Gautier, 2017) implemented in R. Analyses were conducted on SNP datasets both including and excluding transitions to account for possible bias due to DNA degradation. Following the developer’s guidelines, LD pruning was not required, and all runs were performed on unpruned datasets.

A hidden Markov model was built for 12 homozygosity-by-descent (HBD) classes (K) with a rate (R_k_) of 2 and default genotyping and sequencing error rates of 0.001. This error rate was higher than the observed substitution error estimated with ANGSD (see above) to allow for more data flexibility. The use of multiple HBD classes allows a more accurate detection of both short and long HBD segments, resulting in more precise estimates of individual autozygosity (Bertrand et al., 2019). Smaller HBD classes capture longer HBD segments associated with recent ancestry, while larger classes include shorter segments and reflect ancient ancestry. The rate parameter (R_K_ = 2) exponentially increased HBD class age by a factor of 2, allowing finer dissemination between classes. With K set to 12, the model included 11 HBD classes (2, 4, 8, 16, 32, 64, 128, 256, 512, 1024, 2048) and one non-HBD class.

When genetic maps are available, the rate of each HBD class (here, 2–2048) corresponds to approximately twice the number of generations to the most recent common ancestor (Druet & Gautier, 2017). However, here, physical distances between SNPs (in base pairs) were used instead of genetic distances, so the rates could not be directly interpreted in terms of generations (Bertrand et al., 2019). Therefore, HBD segment lengths inferred from the Viterbi algorithm were classified into two length categories for comparison across groups: short to intermediate (1–5 Mb), reflecting more ancient inbreeding, and long (>5 Mb), reflecting recent inbreeding. Segments shorter than 1 Mb were not considered because detection probability decreases in low-coverage data (Tom Druet, personal communication, February 9, 2026).

Individual inbreeding coefficients (F_ROH_) were calculated by summing up the probabilities of HBD states obtained from the forward-backward algorithm for classes 2–128. Although HBD segments identified by the Viterbi algorithm were assigned to classes 16–128 and 2048, F_ROH_ estimation is based on posterior probabilities and therefore includes smaller classes to ensure accurate quantification of recent inbreeding (Tom Druet, personal communication, February 9, 2026). F_ROH_ and HBD length distributions across groups were compared using one-way ANOVA.

### Use of large language models

ChatGPT and Google Gemini were used to assist with command-line and R scripts and to polish the language of the manuscript.

## Results

### Sequencing quality

A total of 26 golden eagles from Finland across three temporal periods (PRE = 7, BOT = 9, POST = 10) were included in whole-genome sequencing analyses. After downsampling to 2× coverage, the breadth of coverage was between 59.0% and 84.0% (Table S1), and the substitution error rate was 0.00065 (Figure S2).

### Drastic loss of genetic diversity

Genetic diversity decreased significantly from PRE to POST across all estimators (Table 1). Median heterozygosity declined by 82% (Figure 2), nucleotide diversity by 42%, and Watterson’s theta by 67%. Differentiation across temporal groups was significant for all estimators (Table S3). The proportion of variable sites also decreased from 0.75% to 0.22%.

**Figure 2.**
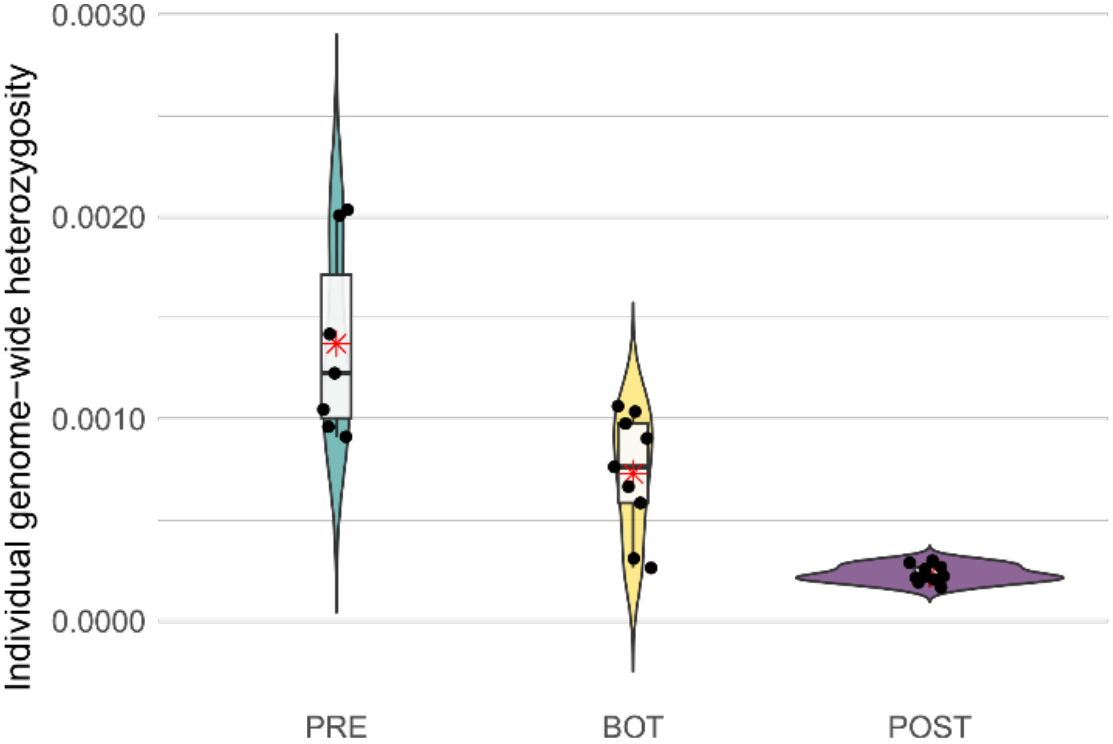
Violin plots of individual genome-wide heterozygosity of Finnish golden eagles across three temporal periods: pre-bottleneck (PRE), bottleneck (BOT), and post-bottleneck (POST). Each black dot represents an individual; the thick blackline within the boxplot indicates the median, and the red asterisk marks the average value.

**Table 1.** Genetic diversity parameters of Finnish golden eagles across three temporal periods: pre-bottleneck (PRE), bottleneck (BOT), and post-bottleneck (POST).

| Parameter | PRE | BOT | POST |
|---|---|---|---|
| Median heterozygosity | 0.0012 | 0.0008 | 0.0002 |
| Nucleotide diversity, π | 0.0019 | 0.0014 | 0.0011 |
| Watterson’s theta, θ<sub>W</sub> | 0.0024 | 0.0015 | 0.0008 |
| Tajima’s D | -1.02 | -0.16 | 1.63 |
| % of variable sites | 0.75 | 0.39 | 0.22 |

### Elevated inbreeding in the modern population and evidence of population bottleneck

The analysis of homozygosity across the genome suggested a trend of increasing inbreeding over time. HBD segment length increased from PRE to POST in both intermediate and long length categories (Table S1, Figure 3). In the 1–5 Mb category, the average total length of HBD segments was 13.7 Mb in PRE, 25.7 Mb in BOT, and 29.0 Mb in POST. For segments >5 Mb, the group means were 4.6, 4.6, and 8.4 Mb, respectively. These averages included individuals without any >5 Mb HBD segments, reducing the average group values to below 5 Mb. The proportion of individuals with at least one segment >5 Mb was 57.1% in PRE, 44.4% in BOT, and 60.0% in POST. Among these individuals, the mean total group length was 8.1 Mb in PRE, 10.3 Mb in BOT, and 14.0 Mb in POST. Individual variation was large, and group differences were not statistically significant (Table S3).

**Figure 3.**
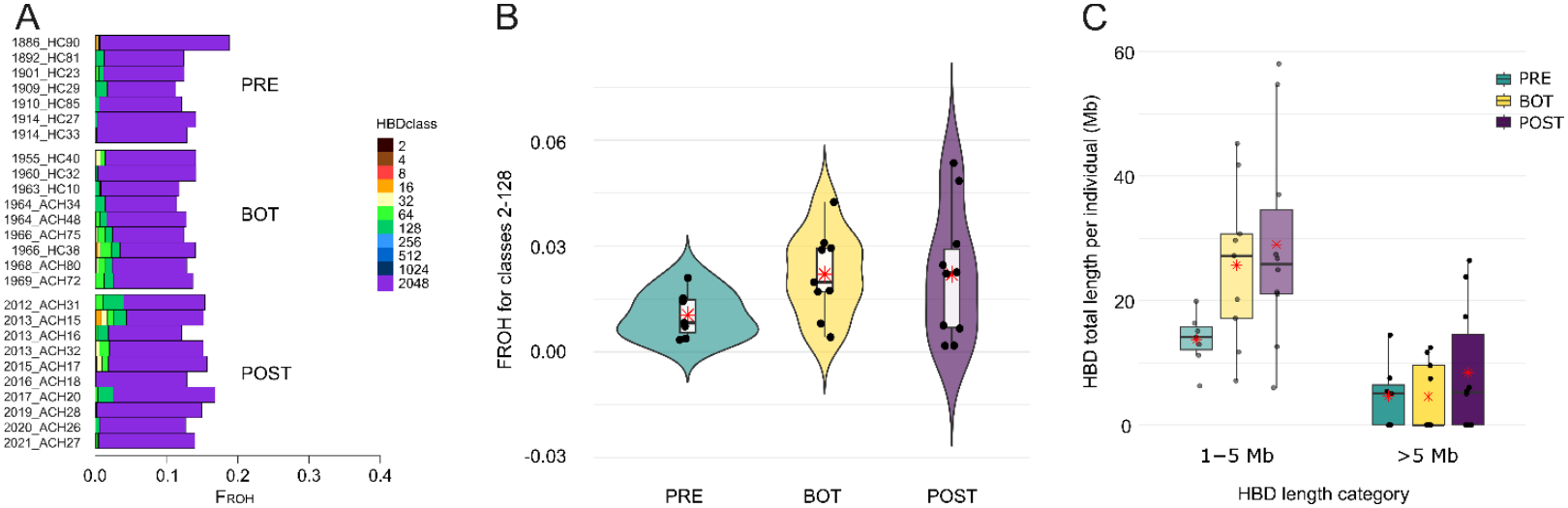
Runs of homozygosity (ROH) analysis of Finnish golden eagles across three temporal periods: pre-bottleneck (PRE), bottleneck (BOT), and post-bottleneck (POST). The analysis is based on 3,149,259 autosomal SNPs, including transitions. **(A)** Homozygous-by-descent (HBD) proportions per class for each individual, ordered chronologically (year_ID). **(B)** Distribution of individual F_ROH_ values (from HBD classes 2–128) across the three groups. Each black dotrepresents an individual, the thick black line within the boxplot indicates the median, and the red asterisk marks the average values. **(C)** Distribution of HBD lengths for the two size categories: 1–5 Mb (intermediate inbreeding; faded color) and >5 Mb (recent inbreeding; bright color). Each black dot represents an individual; the thick black line within the boxplot indicates the median; and red asterisks mark the average values.

Most detected HBD segments were short, associated with ancient inbreeding (Figure 3). Individual genomic inbreeding coefficients (F_ROH_) calculated from recent to intermediate HBD classes ranged from 0.002 to 0.054 (Table S1, Figure 3). Mean F_ROH_ was lowest in PRE (0.010) and higher in BOT and POST (both 0.022), but differences among temporal groups were not statistically significant (Table S3).

A demographic bottleneck was supported by shifts in genome-wide allele frequency spectra. PRE and BOT displayed L-shaped SFS with an excess of low-frequency variants (singletons), particularly prominent in PRE (Figure 4). In contrast, the POST SFS was distorted, showing fewer singletons than other minor allele frequency classes (Figure 4). Even though downsampling most affected the shape of the POST SFS, the pattern of reduced number of low-frequency class minor alleles was observed across all datasets. Furthermore, to validate the observed pattern, we performed an independent downsampling replicate and re-analyzed the SFS metrics. Re-analysis resulted in consistent SFS shapes and fewer singletons in the POST group relative to other minor allele frequency classes.

**Figure 4.**
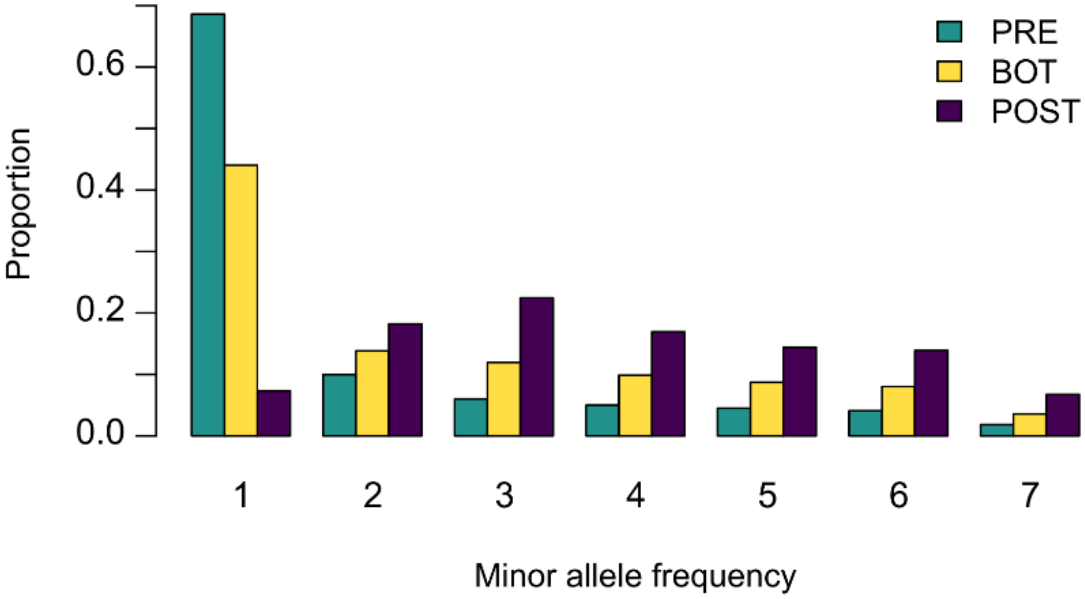
Folded site frequency spectrum (SFS) distributions of variable sites in Finnish golden eagles across three temporal groups: pre-bottleneck (PRE), bottleneck (BOT), and post-bottleneck (POST).

Tajima’s D values shifted gradually from negative in PRE (-1.02) to near-neutral in BOT (-0.16) to positive in POST (1.63) (Table 1), with patterns maintained across the genome rather than being restricted to specific regions (Figure S3); and the across-group differentiation was highly significant (Table S3).

### No clear genetic clustering despite detectable temporal differentiation

After removing transitions and applying LD pruning, the dataset for PCA and Admixture analysis included 22,573 SNPs. Neither PCA nor Admixture analysis revealed clear population structure among PRE, BOT, and POST groups (Figure S4). Admixture analyses supported K = 5 as the most likely number of clusters based on log-likelihood (LnP(K)) and K = 4 based on Evanno’s ΔK (Figure S5), but no visual clustering was observed from the admixture proportions (Figure S4).

Despite the lack of clear clustering, weighted genome-wide pairwise F_ST_ values were 0.043 between PRE and BOT, 0.049 between PRE and POST, and 0.036 between BOT and POST. In all comparisons, the observed F_ST_ values were at or beyond the upper ends of the permutation distributions, indicating that the observed differentiation was representative of the temporal group assignment (Figure S6). Similarly, PBS values were highest for PRE (0.029), followed by POST (0.021), and lowest for BOT (0.015), indicating the highest genetic differentiation of the PRE group.

### Patterns of population differentiation identify putative adaptive loci in the modern population

Manhattan plots of PBS highlighted two genomic regions with elevated differentiation in the POST group (Figure S7). These peaks were not observed in permuted datasets, suggesting that they were associated with the temporal groups rather than random group assignment (Figure S8). One peak was on chromosome 1 from approximately 70.0 to 70.6 Mb – a region overlapping several genes, including protein-coding genes with known cellular regulatory functions (Table S5). A second PBS peak was observed on chromosome 6 from approximately 19.7 to 25.7 Mb, overlapping genes involved in immune functions (Table S5). Meanwhile, PRE and BOT had multiple single-window PBS outliers but did not form clear clusters of genetic divergence in specific genomic regions (Figure S7).

## Discussion

In this study, we used low-coverage whole-genome data spanning 136 years to assess the genetic consequences of the mid-20^th^-century population bottleneck in Finnish golden eagles. Despite a demographically recovering population, we detected a significant loss of genetic diversity, including an 82% reduction in genome-wide heterozygosity, a 42% decline in nucleotide diversity, and a 67% decline in Watterson’s theta. Meanwhile, other results were less straightforward. Although the modern population showed elevated inbreeding, this result was not statistically significant. Similarly, no clear genetic clustering was observed across time, yet pairwise F_ST_ and PBS analyses detected genetic differentiation among temporal groups. These patterns were robust to inclusion/exclusion of transitions and varying sequencing depth, indicating that potential post-mortem DNA damage and historical sample quality did not bias the observed trends.

### Genome-wide signals of the bottleneck

A consistent decline across multiple diversity metrics indicates that the population bottleneck substantially reduced genome-wide genetic variation in golden eagles. The bottleneck was further supported by the observed depletion of rare variants in the modern population, as rare variants typically disappear first during population contractions. In addition, the gradual genome-wide shift in Tajima’s D from negative to positive values over time is also consistent with a demographic transition towards contraction. As expected in bottlenecked populations, inbreeding levels were higher in the modern population, although the increase was not statistically significant, suggesting that the bottleneck did not substantially increase mating among close relatives. This could indicate that the population retained a sufficient number of unrelated individuals or that immigration helped prevent inbreeding. Given that golden eagles in Northern Europe are panmictic and that genetic diversity is low across the region (Karabanina et al., 2024), immigration could have helped maintain relatively low inbreeding without necessarily increasing overall genetic diversity.

Despite the substantial loss of genetic variation, PCA and Admixture analysis did not reveal clear genetic clustering among temporal groups, suggesting that loss of genetic diversity did not lead to the development of distinct genetic groups over the past 100 years. Nevertheless, as evident from the SFS and PBS analyses, the historical population had many unique genetic variants (singletons), which were lost after population contraction. However, further studies are needed to investigate the function of these lost variants to assess their impact on species’ fitness and viability.

When comparing Finnish golden eagles to populations elsewhere in the Holarctic, they show a substantially lower genome-wide heterozygosity (0.0002 in Finland vs. 0.0012–0.0015 across the Holarctic; Sato et al., 2024). Interestingly, nucleotide diversity in Finland (0.0011) is only slightly lower than in other regions (0.0013 and 0.0015, Sato et al., 2024). In contrast, previous studies of golden eagles based on microsatellite markers found similar levels of heterozygosity in Finnish eagles and populations elsewhere across the species’ distribution (Karabanina et al., 2024; Kylmänen et al., 2023; Nebel et al., 2015, 2023). This discrepancy may partly reflect differences in marker types. Whole-genome data capture variation at thousands of SNPs distributed throughout the genome, including both coding and non-coding regions, whereas microsatellite studies usually use a limited number of neutral loci. Furthermore, microsatellites generally have higher allelic diversity per locus than mostly biallelic SNPs, and in population genetics, including the study of Finnish golden eagles by Kylmänen et al. (2023), only the most polymorphic microsatellites are used, which may have inflated genetic diversity estimates. Thus, as shown by Väli et al. (2008), populations having the same level of microsatellite heterozygosity may have very different levels of genomic diversity.

### Demographic recovery vs. genetic erosion

The Finnish golden eagle population has grown 40–50% since the early 2000s, with the number of breeding pairs increasing from 250–300 to 365–469 by 2025 (Koskimies & Ollila, 2024; Sundvall, 2025). However, reproductive output has slowly declined over the last decades, from a long-term average of 0.53 chicks per occupied territory since the beginning of monitoring in the 1970s to 0.37 in 2025 (Sundvall, 2025). While the causes for this decline remain unknown, low genetic diversity or loss of genetic variants following the mid-20^th^ century bottleneck may have caused a higher genetic load that partly limits fitness and reproductive success (e.g., Heber and Briskie, 2010). Here, we identified several regions of elevated population differentiation in the modern population (PBS peaks) overlapping with protein-coding and regulatory genes. However, the functional significance of these regions and their potential effects on genetic load and fitness remain unknown. Therefore, further research is needed to determine whether genetic changes following the bottleneck have contributed to the decline in reproductive output.

When compared to other birds that have undergone similar human-induced declines and subsequent recoveries, the heterozygosity level in Finnish golden eagles is alarmingly low. Their genome-wide heterozygosity (0.0002) is lower than that of peregrine falcons (*Falco peregrinus*; 0.0010), the bald and the white-tailed eagles (*Haliaeetus leucocephalus*, *H. albicilla*, both 0.0004), Philippine eagles (*Pithecophaga jefferyi*, 0.0003), and even the critically endangered Kākāpōs (*Strigops habroptilus*, 0.0005–0.0009) (Dussex et al., 2021; S. Li et al., 2014; Perdon et al., 2025). Although direct numerical comparisons are not possible due to methodological differences among studies, these estimates suggest that despite demographic recovery, Finnish golden eagles may be more genetically depleted than many species currently listed as threatened. This emphasizes an important conservation point: increasing the number of individuals does not automatically restore lost genetic variation (Frankham et al., 2010; Lande et al., 2010).

### Demographic history

The census size of the Finnish golden eagle population before the mid-20^th^ century bottleneck is unknown, but the 40% decline in nucleotide diversity and 80% decline in heterozygosity suggest a considerably larger population prior to the bottleneck, particularly because long generation times can buffer the loss of genetic diversity (e.g., Hailer et al., 2006). According to Tuomo Ollila (personal communication, October 23, 2026), who was responsible for golden eagle monitoring in Finland for two decades at Parks and Wildlife Finland (Metsähallitus), the population size before the bottleneck could have been at least 700–1000 breeding pairs based on habitat availability and land use. However, our pre-bottleneck group likely does not represent the true historical baseline. Historical records state that persecution of large raptors had already intensified by the 18^th^ and 19^th^ centuries (Erkamo, 1990; Vuorisalo & Laihonen, 2000), suggesting that population decline may have begun before the earliest samples used here. Consequently, the genetic diversity observed in the pre-bottleneck group may already reflect a partial population decline. Interestingly, the excess of singletons in this PRE group’s SFS suggests recent population growth, potentially indicating demographic fluctuations preceding the bottleneck, although limited temporal sampling from that period does not allow for making definitive conclusions.

Runs of homozygosity (ROH) analyses further suggested that the mid-20^th^ century bottleneck in Finnish golden eagles occurred in a population that had already been shaped by ancient demographic constraints. Most homozygous-by-descent (HBD) segments belonged to the oldest age class, indicating ancient inbreeding, likely associated with post-glacial recolonization of Fennoscandia after the Scandinavian Ice Sheet retreat (10–12 ka; Bertrand et al., 2019; Regnéll et al., 2023).

### Conservation implications

Preserving genetic diversity is a core component of biodiversity, as recognized by the Kunming-Montreal Global Biodiversity Framework (United Nations Convention on Biological Diversity, 2022). Our results indicate that management decisions based solely on census growth risk overlooking reduced long-term resilience of species. For example, in Finland, golden eagles have already shown reduced reproductive output over the last decades (see above), which, combined with findings of this study, may reflect reduced genetic health following population contraction during the past centuries.

Furthermore, although the golden eagle is globally listed as Least Concern due to its large range and stable population size (BirdLife International, 2023), our study emphasizes the need for regional population assessments to inform conservation, particularly in regions with documented historical bottlenecks and when species show large-scale population differentiation (Karabanina et al., 2024; Nebel et al., 2015; Sato et al., 2020).

At local levels, including in Finland, golden eagles continue to face multiple pressures, including human disturbance, habitat loss, collisions with vehicles and infrastructure, and accelerated climate change in the Arctic (four times faster than in lower latitudes) (Ollila, 2018; Rantanen et al., 2022). To support the long-term viability of the population, conservation actions should promote both demographic stability and the preservation of genetic diversity and adaptive potential. In addition to maintaining legal protection, management efforts should prioritize increasing breeding success to support population recovery and reducing human-caused mortality to prevent further declines. Facilitating gene flow between populations could help restore genetic variation lost during the bottleneck and prevent inbreeding. Altogether, these actions would support the long-term genetic health and adaptive potential of golden eagles not only in Finland but in the whole North European region.

## Acknowledgements

We thank Soile Alatalo for laboratory assistance; Tuula Pudas for help with the museum collections at the Oulu Zoological Museum; and Torsten Stjernberg, Martti Hildén, and Roni Andersson for assistance with the collections at the Finnish Museum of Natural History. We also thank Aure Kylmänen and Tobias Rapsch for their help with museum sampling. We are grateful to Tom Druet for advice on the RZooROH analyses. We thank Juan Galarza, Samuel Lam, and Stefan Prost for advice on bioinformatic analyses, Saskia Wutke for valuable comments on the manuscript, and Heikki M. Salo for artistic and writing advice. This work was funded by the Kvantum Institute at the University of Oulu, the University of Oulu Scholarship Foundation, and Interreg Aurora. We also acknowledge CSC – IT Center for Science, Finland, for providing computational resources, and Novogene for sequencing services.

## Author contribution statement

EK and LK led the study design. EK and SP performed the sampling. EK conducted the lab work, performed data analyses, generated scripts, figures, and tables, and drafted the manuscript. SP and TM contributed to data validation. AN contributed to data analysis. LK, TM, SP, and AN supervised the work. LK and EK acquired funding. All authors revised the manuscript.

## Data archiving

Raw whole-genome sequencing read data for the 27 golden eagles sequenced in this study are openly available at the European Nucleotide Archive under project accession PRJEB120806. The scripts of the bioinformatics analyses are available on GitHub under “lcWGS-genomic-erosion-pipeline”: https://github.com/ekarabanina/lcWGS-genomic-erosion-pipeline/tree/7e395d87d641f15ae28fe0fa241fce269e85b996.

## Research ethics statement

This study used samples of golden eagles obtained from Finnish museum collections. The species is protected in Finland, and all specimens were originally collected and curated by the museums under national legislation. Access to the material was granted by the representatives of the museums. No ethical approval was required. No animals were captured or harmed for this research.

