## Supplementary information for "Genome-wide diversity across 130 years reveals drastic post-bottleneck genetic erosion in a recovering population of golden eagles *(Aquila chrysaetos)*"

#### Excel tables ("Suppelmentary_tables_S1-S5"):

1. **Table S1:** Sample information, lab and sequencing results, heterozygosity, and summary of HBD segments from Viterbi algorithm.
2. **Table S2:** Overview of analyses performed for each dataset.
3. **Table S3:** Comparison of results from different datasets.
4. **Table S4:** Probabilities of HBD segments from forward-backward algorithm for each class for the three datasets: (1) downsampled to 2x including transitions, (2) downsampled to 2x excluding transitions, (3) full depth including transitions.
5. **Table S5:** List of all genes overlapping PBS peaks on chromosomes 1 and 6 in the modern population.

#### PDF file ("Supplementary_information"):

1. **Appendix 1:** Phenol-chloroform DNA extraction protocol.
2. **Appendix 2:** Block scheme of bioinformatics pipeline and a link to GitHub scripts.
3. **Appendix 3:** Results.

**3.1 Primary dataset (2x with transitions), additional results**

- Figure S1: LD-decay plot (only transversions)
- Figure S2: Substitution error rate
- Figure S3: Manhattan plots of genome-wide Tajima's D
- Figure S4: PCA plot and Admixture proportions
- Figure S5: Admixture results: LnP(K) and Evanno's ∆K plots
- Figure S6: Distribution of permutation values of $F_{ST}$
- Figure S7: Manhattan plots of PBS values
- Figure S8: Distribution of permutation values of PBS

**3.2 LD pruning, PCA, and Admixture (full depth)**

- Figure S9: LD-decay plot (only transversions)
- Figure S10: PCA plot
- Figure S11: Admixture results: LnP(K), Evanno's ∆K, and admixture proportions plots

**3.3 Effect of excluding transitions and uneven sequencing depth**

- Figure S12: Substitution error rate (full depth)
- Figure S13: Individual genome-wide heterozygosity across groups
- Figure S14: Correlation between heterozygosity and sequencing depth
- Figure S15: Manhattan plots of PBS
- Figure S16: Runs of homozygosity (ROH): HBD proportions and $F_{ROH}$ across groups
- Figure S17: Distribution of HBD lengths for the two size categories
- Figure S18: Correlation of ROH parameters with sequencing depth
- Figure S19: SFS plots

### APPENDIX 1: PHENOL-CHLOROFORM DNA EXTRACTION PROTOCOL

**Clean laboratory:**

A clean laboratory at the University of Oulu (Oulu, Finland) is dedicated to working with historical or otherwise sensitive samples. It is UV-treated twice a day for 2 hours each time. Surfaces, scissors, tweezers, and scalpels were sterilized with 10% bleach followed by 70% ethanol. All work involving phenol and chloroform was done in a fume hood.

**Reagents:**

- STE buffer, pH = 8.0
    - 0.1 M NaCl
    - 10 mM Tris-HCl
    - 0.1 mM EDTA
- SET buffer, pH = 8.0
    - 0.15 M NaCl
    - 1 mM EDTA
    - 50 mM Maleic acid
- 10% SDS (Sodium Dodecyl Sulfate)
- EB buffer (10 mM Tris-HCl, pH = 8.5)
- Phenol
- Chloroform
- Phenol-chloroform 1:1
- 3 M sodium acetate
- Proteinase K (20 mg/ml)
- 1 M DTT (dithiothreitol)
- Absolute ethanol
- 70% ethanol
- 10% bleach
- Sterile $H_2O$

**Washing the samples (toe pads, skins):**

Samples were rinsed with 10% bleach, then with 96% ethanol and finally several times with sterile water. The tube was changed before the last water wash.

**Rehydrating the samples (muscle tissue in ethanol, toe pads, skins):**

Samples were rehydrated with STE buffer for 5 min at room temperature.

**DNA extraction protocol:**

1. Cut the tissue into tiny pieces and place them into the Eppendorf tube.
2. Add:
    - 400 µl SET buffer
    - 14 µl 10% SDS
    - 10 µl proteinase K (20 mg/ml)

- 10 µl 1M DTT to skin and tissue samples; 20 µl 1M DTT to toe pad samples or to skins with lots of feathers

3. Mix carefully by pipetting.
4. Incubate overnight at 55°C.
5. Vortex to mix thoroughly.
6. Spin to settle down the undissolved particles (if needed).
7. Transfer the liquid phase to the new tube, avoiding any solid undissolved particles.
8. Add 400 µl of saturated phenol, vortex, and centrifuge 5 min at 13 000 rpm.
9. Transfer the upper phase (contains DNA) into a new tube, add 400 µl of phenol-chloroform (1:1), vortex and centrifuge 5 min at maximum speed (13,000 rpm).
10. Transfer the upper phase into a new tube, add 400 µl of chloroform, vortex, and centrifuge **2 min** at 13,000 rpm.
11. Transfer the upper phase into a new tube, add 2x volume of ice-cold absolute ethanol (99.5%), mix carefully by turning the tube upside-down several times.
12. Keep at -20 °C overnight to enhance the precipitation of the DNA.
13. Centrifuge 30 min at 13,000 rpm.
14. Pre-heat EB buffer at 50°C.
15. Remove the ethanol carefully (keep the DNA pellet at the bottom of the tube).
16. Wash with 70% ethanol: add 200 µl ethanol, DO NOT MIX, centrifuge 5 min at 13,000 rpm.
17. Remove the ethanol. Keep the DNA pellet untouched in the bottom of the tube.
18. Dry the pellet by keeping the caps open at room temperature for about an hour OR in a heat-block at 65°C for a few minutes.
19. Dissolve into 100 µl EB buffer (warm the EB buffer prior dilution). If the pellet is very small or invisible, dilute into 50 µl of EB buffer. Mix carefully by pipetting.
20. Measure the DNA concentration and purity (if needed, use EB buffer as blank).
21. Store DNA extracts at -20 °C.

### APPENDIX 2: BIOINFORMATICS PIPELINE

Scripts are provided on GitHub: https://github.com/ekarabanina/lcWGS-genomic-erosion-pipeline

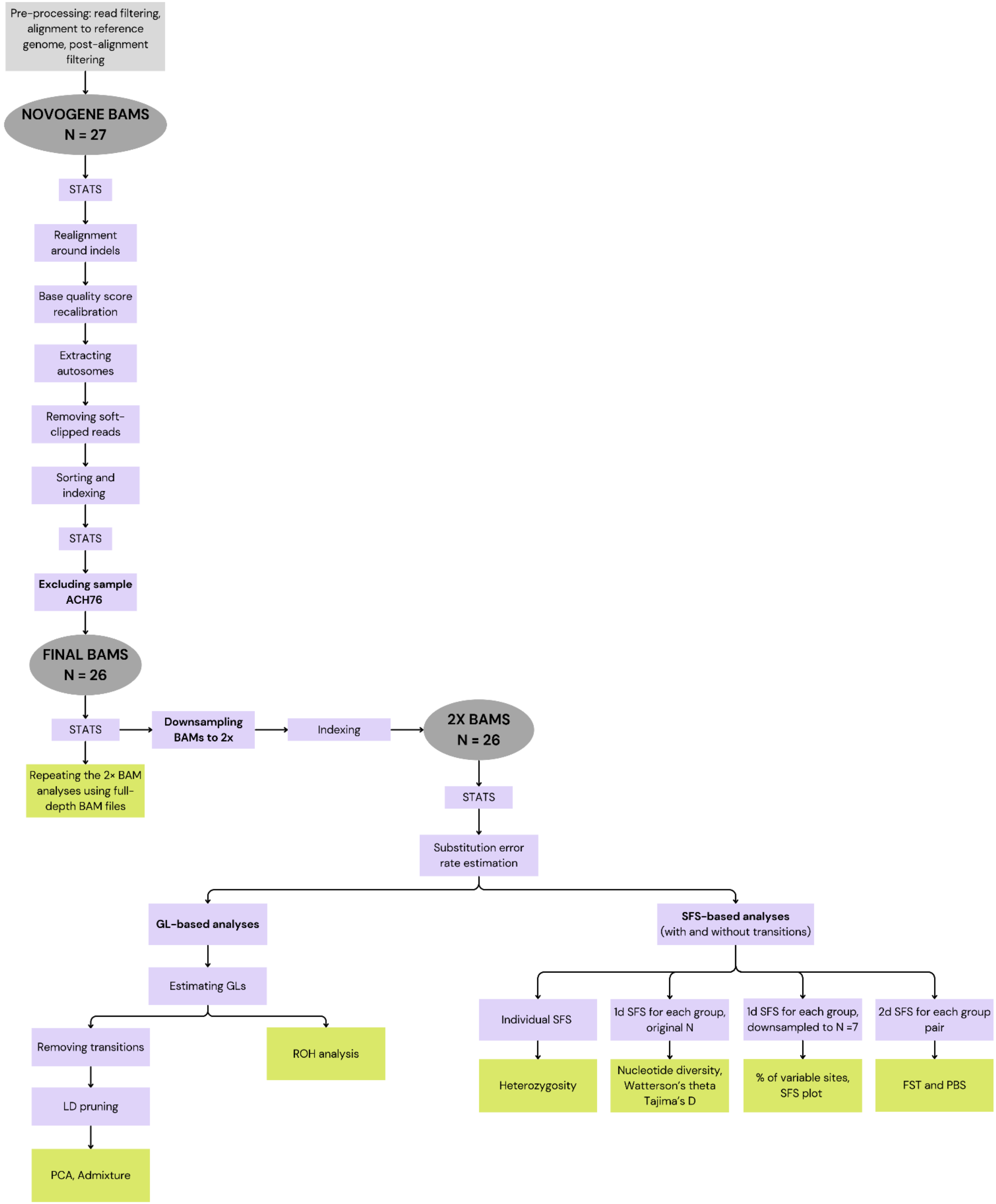

### APPENDIX 3: RESULTS

#### 3.1 PRIMARY DATASET (2X WITH TRANSITIONS), ADDITIONAL RESULTS

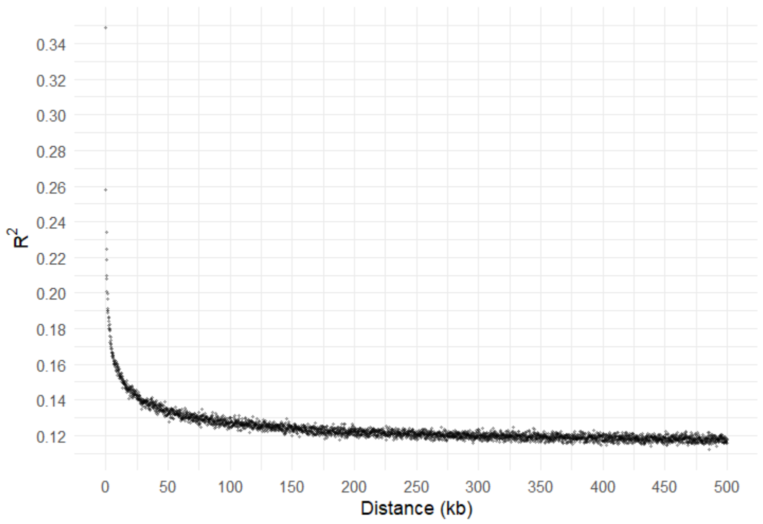


Figure S1. A linkage disequilibrium (LD) decay plot **for transversion SNPs** using ngsLD. Each dot shows an LD score ($R^2$) for each SNP pair within 500 kb in a genome, and a decay curve was generated from a random 10% subset of the data binned into 150 distance intervals.

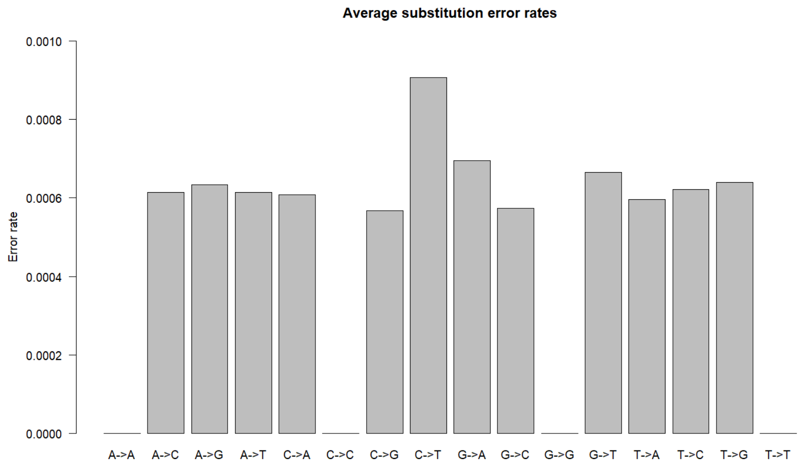


Figure S2. Distribution of substitution error rates for 26 Finnish golden eagles (**2x dataset, with transitions**), estimated per site from genotype likelihoods using ANGSD.

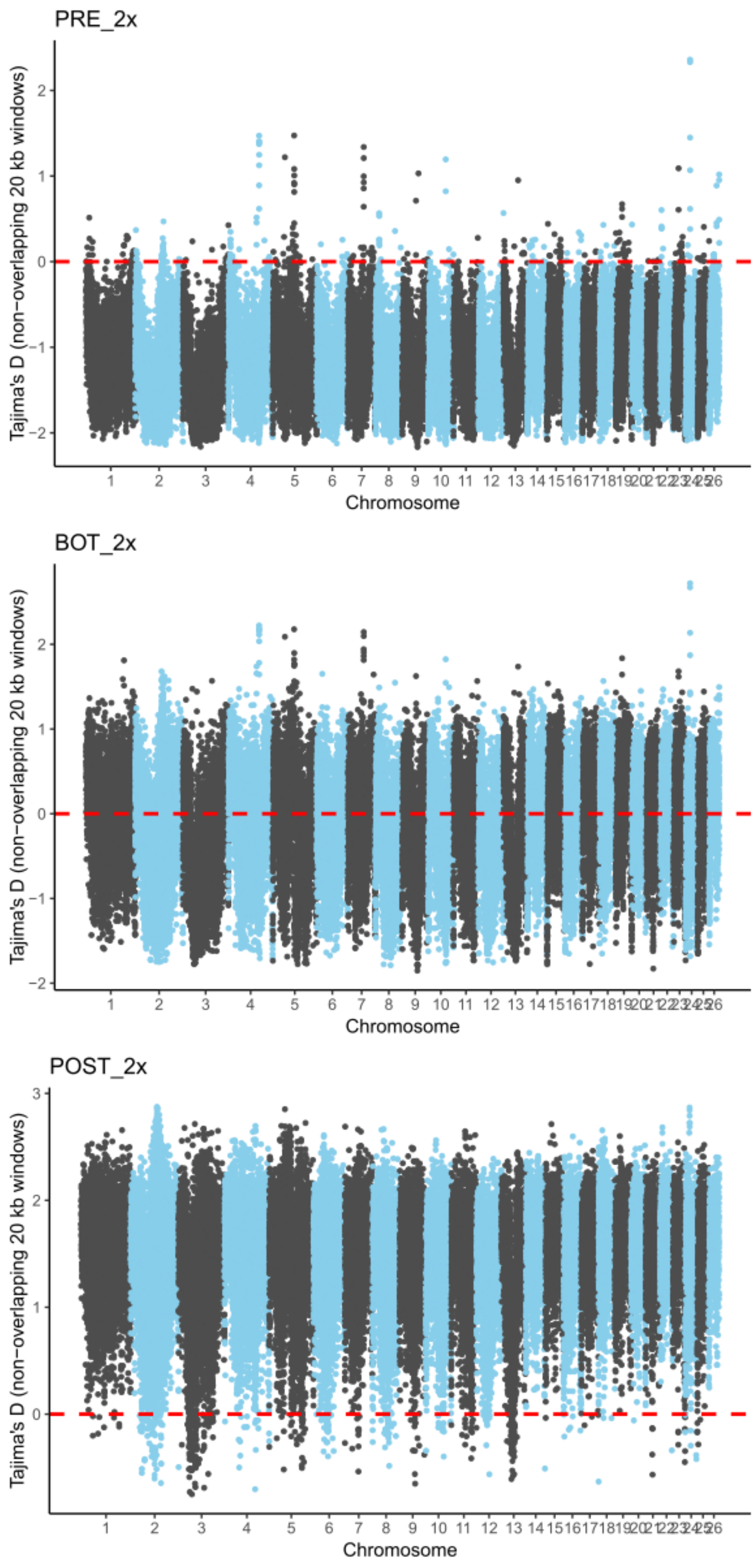


Figure S3. Manhattan plots of genome-wide Tajima's D values of Finnish golden eagles across three temporal periods: pre-bottleneck (PRE), bottleneck (BOT), and post-bottleneck (POST). Each dot is a Tajima's D value for a 20 kbp non-overlapping window in 26 autosomes. Note the different Y-axis scales. The red line indicates Tajima's D of 0.

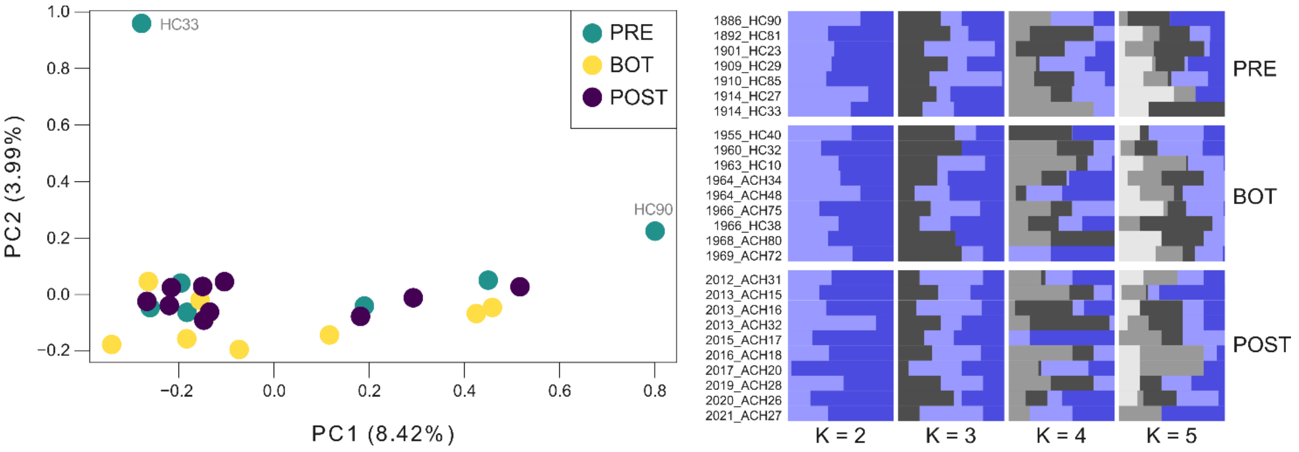


Figure S4. PCA and admixture analyses of Finnish golden eagles across three temporal periods: pre-bottleneck (PRE), bottleneck (BOT), and post-bottleneck (POST). Analyses were based on 22,573 transversion SNPs. **(a)** Principal Component Analysis (PCA) showing that the total variance of 28.46% was best explained by the first principal component (PC1) according to the Minimum Average Partial (MAP) test. The subsequent PCs explained approximately 4% each, with the last PC explaining only 1% of variance. Outliers are complemented with individual IDs. Eigenvectors are scaled according to the proportion of variance explained. **(b)** Admixture analysis showing individual ancestry proportions for K = 2 to 5, with individuals ordered chronologically within each temporal group and labelled as "year_ID".

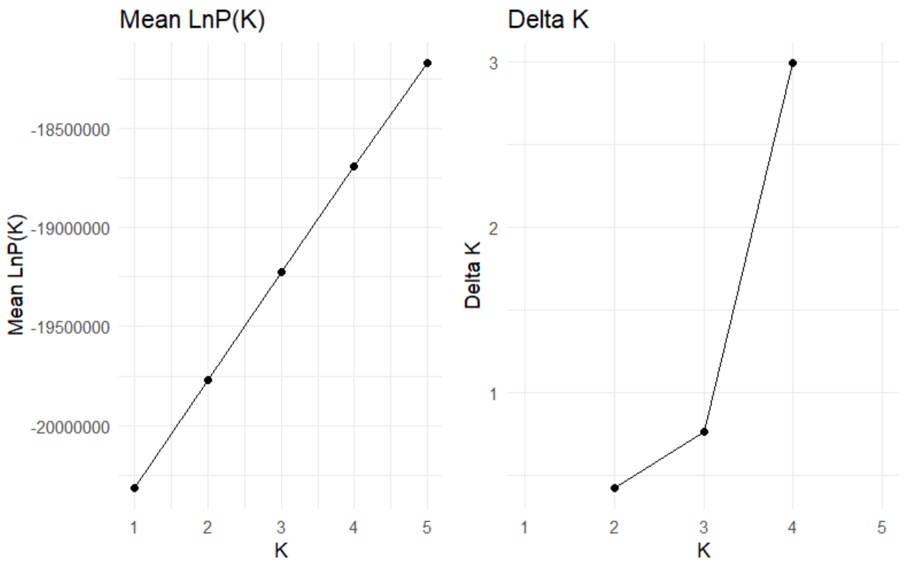


Figure S5. Admixture results for 26 Finnish golden eagles (**2× transversions-only, LD-pruned dataset**) tested for K = 1–5. The left panel shows LnP(K), and the right panel shows Evanno's ΔK. Higher values indicate stronger support for the corresponding K.

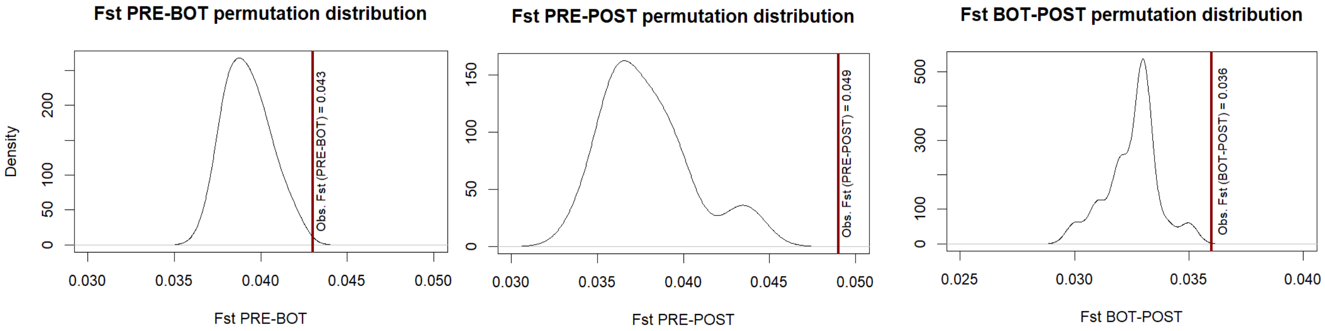


Figure S6. Distribution of weighted pairwise $F_{ST}$ values across 20 permutations for each pairwise comparison (PRE–BOT, PRE–POST, BOT–POST). The observed $F_{ST}$ values are indicated by vertical red lines. Note the different axis scales.

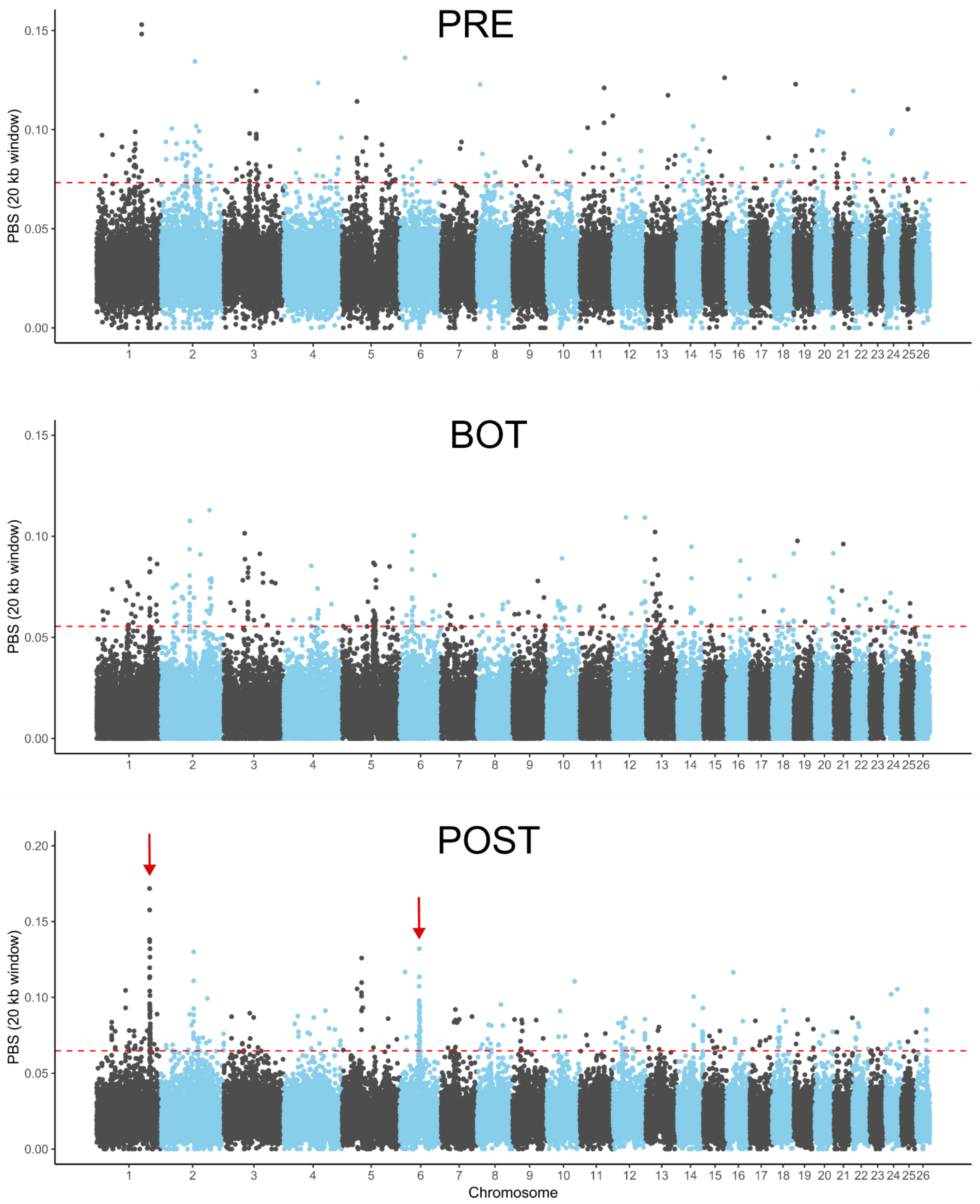


Figure S7. Manhattan plots of population branch statistics (PBS) of Finnish golden eagles across three temporal periods: pre-bottleneck (PRE), bottleneck (BOT), and post-bottleneck (POST). Each dot is a PBS value for a 20 kpb non-overlapping window in 26 autosomes. Note the different Y-axis scales. Red dashed lines show PBS levels equal to four standard deviations above the average for each group. Red arrows point to peaks of differentiation at chromosomes 1 and 6.

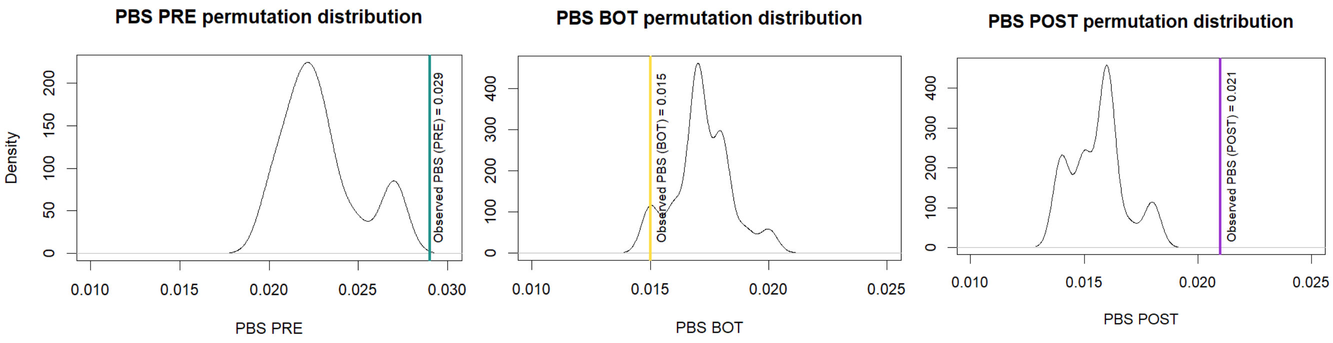


Figure S8. Distribution of weighted PBS values across 20 permutations for each group: pre-bottleneck (PRE), bottleneck (BOT), and post-bottleneck (POST). The observed PBS values are indicated by vertical lines. Note the different axis scales.

#### 3.2 LD PRUNING, PCA, AND ADMIXTURE (FULL-DEPTH)

For the full-depth dataset, principal component analysis (PCA) and Admixture analysis were performed on a linkage disequilibrium (LD)-pruned set of transversion SNPs. Pruning was based on the LD-decay curve (Figure S9) for locus pairs within 500 kb with $R^2 > 0.7$. As in the downsampled dataset, neither PCA nor Admixture showed clear clustering among temporal groups (Figures S10, S11). The most likely number of clusters was K = 5 based on log-likelihood (LnP(K)) and K = 2 based on Evanno's ΔK (Figure S11). One notable difference between the datasets was the group that the outlier individuals were from in PCA: in the downsampled dataset, outliers were from the PRE group, whereas in the full-depth dataset, they were from the POST group.

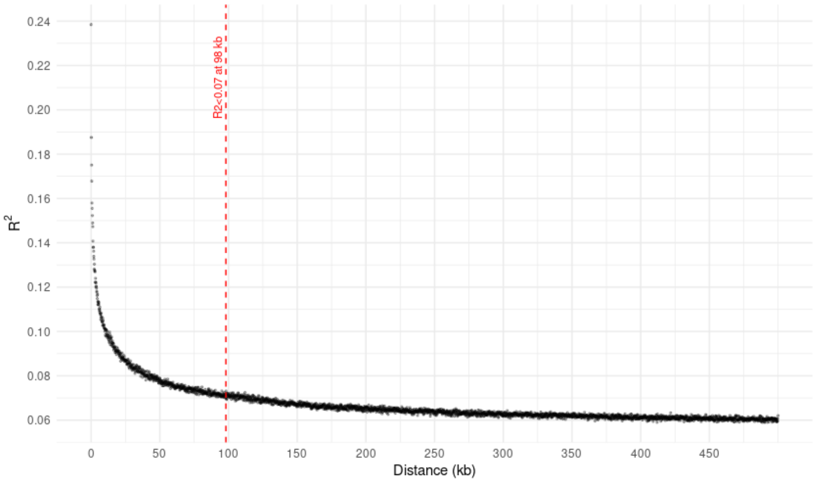


Figure S9. A linkage disequilibrium (LD) decay plot for the total **full-depth dataset** of Finnish golden eagles **using only transversion SNPs**.

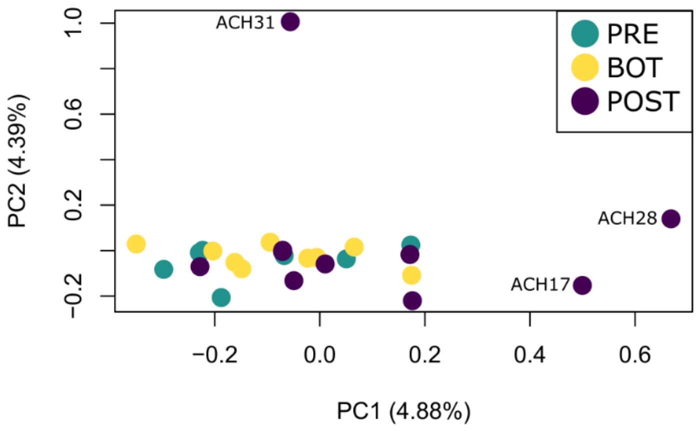


Figure S10. Principal Component Analysis (PCA) of Finnish golden eagles from three temporal periods: pre-bottleneck (PRE), bottleneck (BOT), and post-bottleneck (POST). Analyses were based on **32,351 LD-pruned transversion SNPs**. The total variance was 27.5%. Eigenvectors were scaled according to the proportion of variance explained.

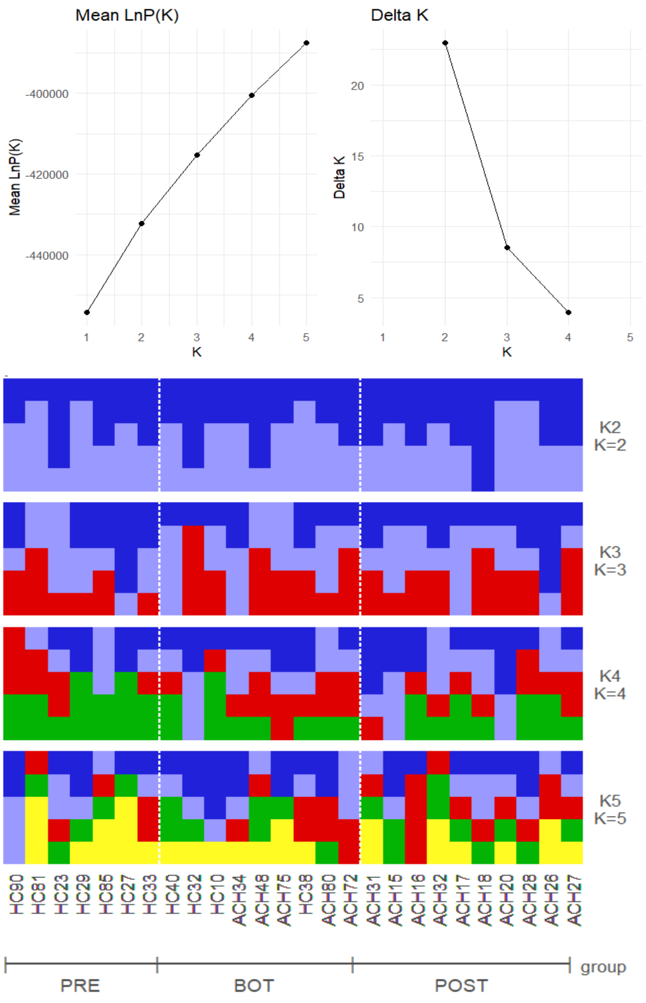


Figure S11. Admixture results for 26 Finnish golden eagles from three temporal periods: pre-bottleneck (PRE), bottleneck (BOT), and post-bottleneck (POST), for the **full-depth, transversions-only, LD-pruned dataset** tested for K = 1–5. The top left panel shows LnP(K), and the top right panel shows Evanno's ΔK. Higher values indicate stronger support for the corresponding K. The bottom panel shows individual ancestry proportions for K = 2 to 5, with individuals ordered chronologically within each temporal group.

#### 3.3 EFFECT OF EXCLUDING TRANSITIONS AND UNEVEN SEQUENCING DEPTH

To evaluate the robustness of diversity estimates to data filtering and sequencing depth, we compared the main dataset (downsampled to 2× with transitions) with two supplementary datasets: (1) downsampled to 2× excluding transitions and (2) the full-depth dataset including transitions (Table S2). Across all datasets, temporal patterns were highly consistent, supporting a robust signal of diversity decline and population contraction.

##### SEQUENCING QUALITY AND DNA DAMAGE

In the full-depth dataset, individual mean sequencing depth ranged from 2.08× to 9.73×, with breadth of coverage between 66.4% and 99.4% (Table S1). The average substitution error rate was low (0.00078; Figure S12). MapDamage2 profiles showed no detectable fragmentation or misincorporation patterns (not presented), indicating minimal post-mortem damage and supporting the inclusion of transition sites in the main analyses.

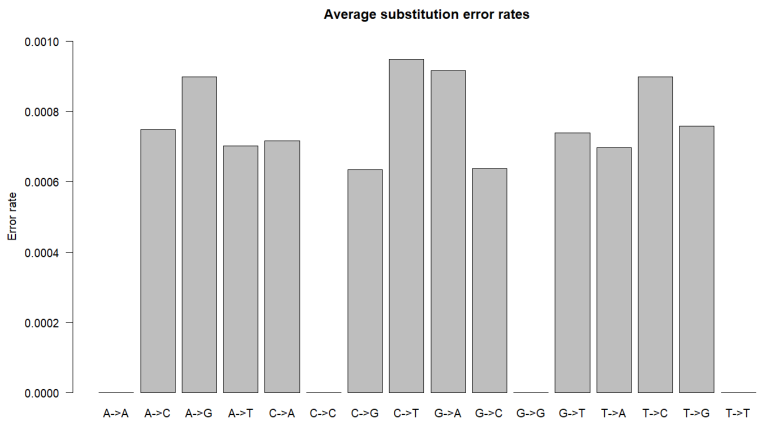


Figure S12. Distribution of substitution error rates for 26 Finnish golden eagles (**full-depth dataset with transitions**), estimated per site from genotype likelihoods using ANGSD.

##### GENETIC DIVERSITY

Across all datasets, genome-wide heterozygosity, nucleotide diversity (π), Watterson's theta ($\theta_W$), and the proportion of variable sites showed a decline of genetic diversity from PRE to POST, with significant temporal differentiation among temporal groups (Table S3, Figure S13). Pairwise comparisons in heterozygosity consistently identified POST as significantly reduced relative to PRE and BOT, whereas PRE and BOT did not differ significantly based on transversion sites only (Table S3).

Excluding transitions reduced absolute genetic diversity values, as expected, but did not change the temporal pattern (Table S3). Similarly, estimates from the full-depth dataset closely matched those from the downsampled data, although the magnitude of heterozygosity decline was smaller.

Individual heterozygosity showed a strong negative correlation with average sequencing depth (Spearman's $\rho$ = -0.76, p = $1.0 \times 10^{-5}$; Figure S14), and therefore the main analyses were conducted using the downsampled dataset to account for differences in sequencing depth between modern and historical samples. However, lower heterozygosity in the POST group remained after downsampling, indicating that the observed reduction in the full-depth dataset was caused solely by sequencing depth.

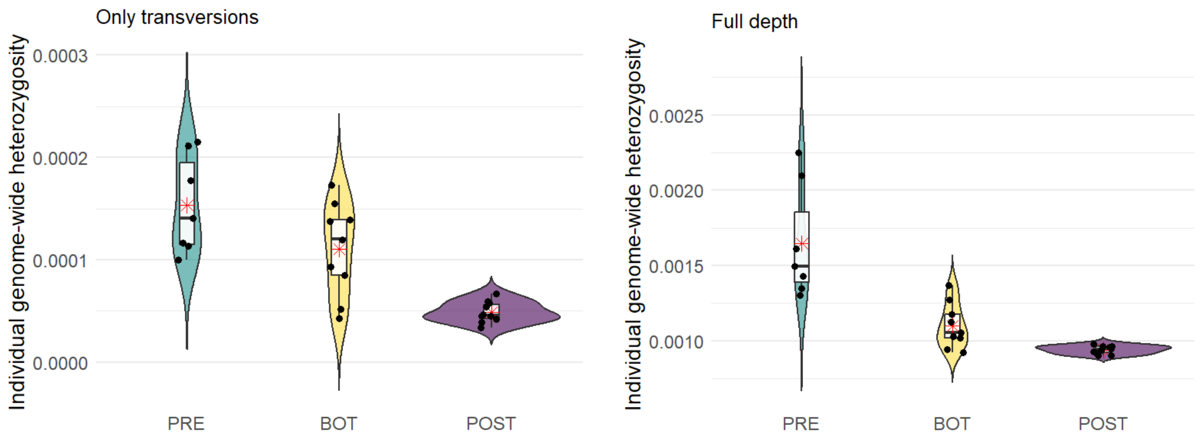


Figure S13. Violin plots showing group-level comparisons of individual genome-wide heterozygosity of Finnish golden eagles across three temporal periods: pre-bottleneck (PRE), bottleneck (BOT), and post-bottleneck (POST) **based on the downsampled dataset without transitions (left) and full-depth dataset with transitions (right)**. Each black dot represents an individual; the thick black line within the boxplot indicates the median, and red asterisks mark the means. Note different Y-axis scales.

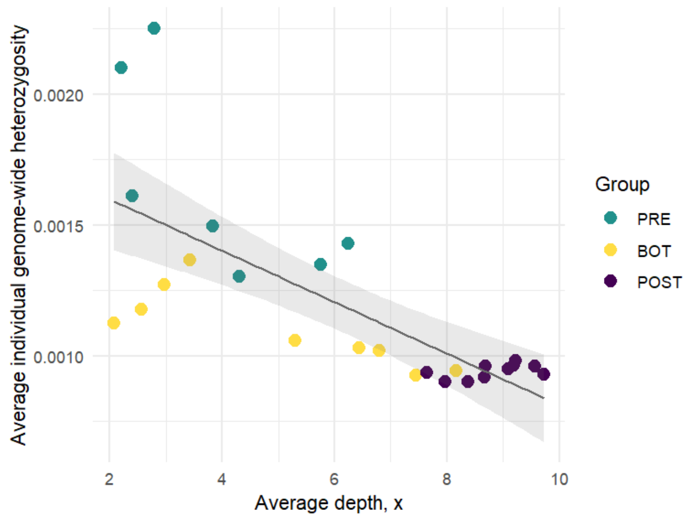


Figure S14. Correlation of individual genome-wide heterozygosity with average sequencing depth for Finnish golden eagles from three temporal groups: pre-bottleneck (PRE), bottleneck (BOT), and post-bottleneck (POST). Each dot is an individual, and colors correspond to temporal groups. The grey solid line is a linear trendline, and the shaded areas are 95% confidence intervals.

#### $F_{ST}$ AND PBS

Population differentiation patterns were consistent across datasets, with PRE showing the strongest differentiation relative to BOT and POST (Table S3). Excluding transitions or using full-depth data resulted in minor differences in absolute $F_{ST}$ and PBS values but did not affect the relative ranking among temporal groups. Manhattan plots of PBS revealed peaks in the same genomic regions across datasets, including shared outlier regions in the POST group on chromosomes 1 and 6 (Figure S15). Although the magnitude of differentiation (i.e., number of outliers in the same region) varied slightly, their genomic positions were consistent.

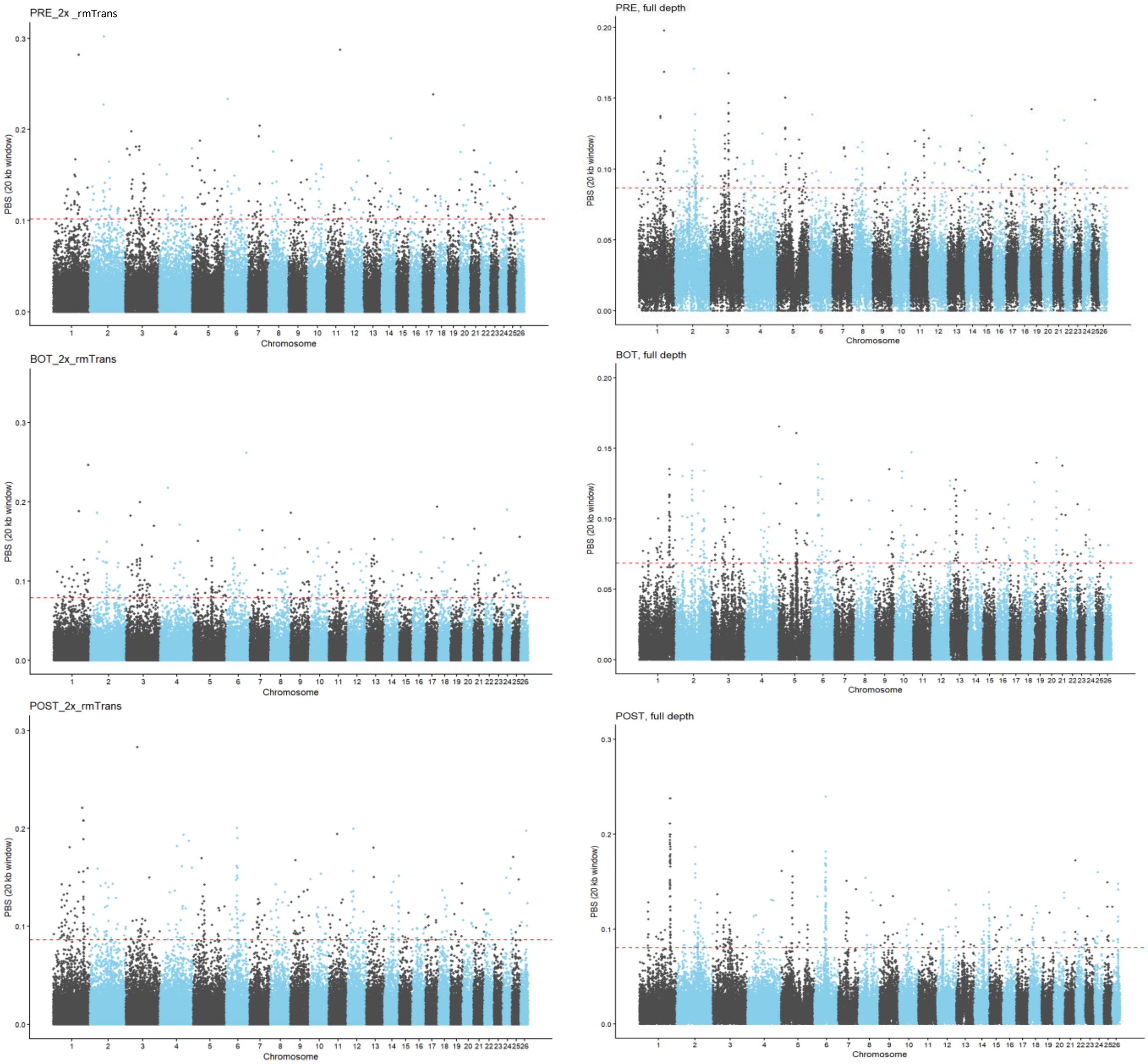


Figure S15. Manhattan plots of population branch statistics (PBS) of Finnish golden eagles across three temporal periods: pre-bottleneck (PRE), bottleneck (BOT), and post-bottleneck (POST) **based on the downsampled dataset without transitions (left) and full-depth dataset with transitions (right)**. Each dot is a PBS value for a 20 kpb non-overlapping window in 26 autosomes. Red dashed lines show PBS levels equal to four standard deviations above the average for each group. Note the different Y-axis scales between the datasets and of the POST group in the full-depth dataset. Red dashed lines show PBS levels equal to four standard deviations above the average for each group.

#### INBREEDING AND RUNS OF HOMOZYGOSITY

Runs of homozygosity (ROH) analyses showed dataset-dependent patterns. Trends from the two downsampled datasets (with and without transitions) were consistent and indicated increasing recent inbreeding toward the POST group (Table S3, Figures S16, S17). In addition, across all datasets, individual inbreeding coefficients ($F_{ROH}$) and total HBD length in the 1–5 Mb category were highest in the POST group (Table S3, Figure S17). However, the full-depth dataset showed an opposite temporal pattern for long HBD segments (>5 Mb), which decreased from PRE to POST (Table S3, Figure S17).

Given that ROH inference is sensitive to uneven coverage in low-depth genomes (Forneris et al., in preparation), the downsampled results likely provide the more reliable signal and suggest a moderate increase in recent inbreeding in the modern population. Furthermore, in the full-depth data, the average HBD segment length and $F_{ROH}$ were strongly correlated with sequencing depth (Figure S18). HBD segment length was negatively correlated (Pearson's r = -0.87, p = $6.2 \times 10^{-9}$), whereas $F_{ROH}$ was positively correlated (Pearson's r = 0.65, p = $3.4 \times 10^{-4}$).

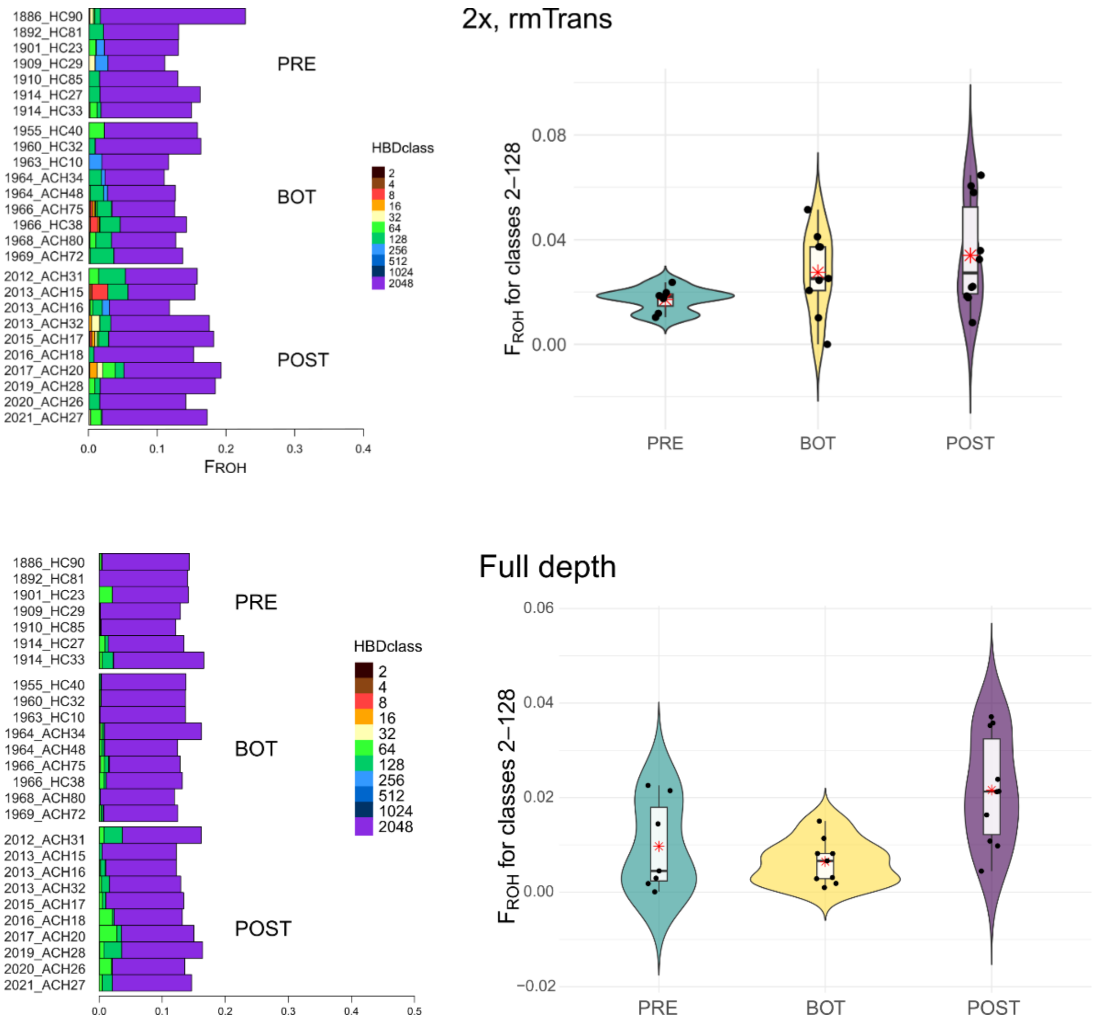


Figure S16. Runs of homozygosity (ROH) analysis of Finnish golden eagles across three temporal periods: pre-bottleneck (PRE), bottleneck (BOT), and post-bottleneck (POST) **based on the downsampled dataset without transitions (846,418 transversion SNPs; top) and full-depth dataset with transitions (4,816,564 SNPs; bottom). (left)** Homozygous-by-descent (HBD) proportions per class for each individual, ordered chronologically (year_ID). **(right)** Distribution of individual $F_{ROH}$ values (from HBD classes 2–128) across the three groups. Each black dot represents an individual, the thick black line within the boxplot indicates the median, and red asterisks mark the average values.

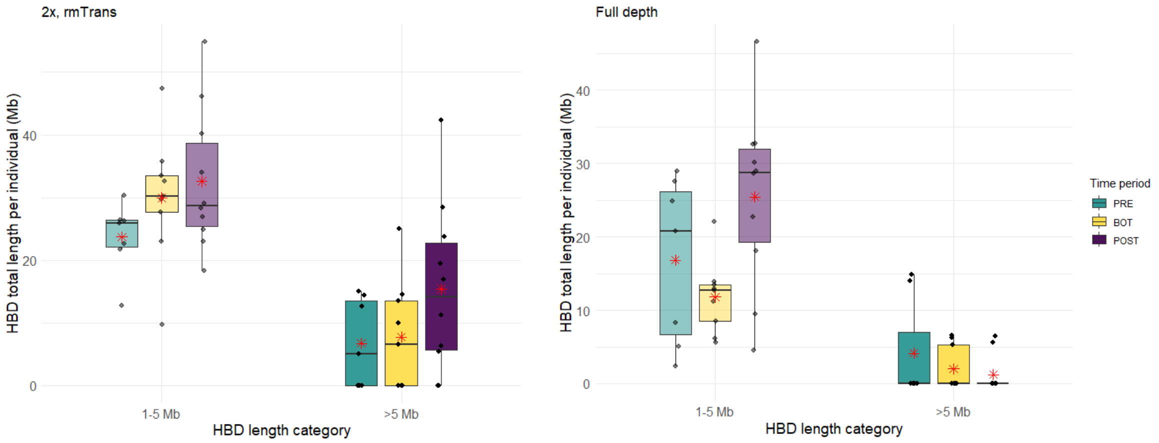


Figure S17. Distribution of HBD lengths for the two size categories: 1–5 Mb (intermediate inbreeding) and >5 Mb (recent inbreeding). The analysis is **based on the downsampled dataset without transitions (846,418 transversion SNPs; left) and full-depth dataset with transitions (4,816,564 SNPs; bottom)**Each black dot represents an individual, the thick black line within the boxplot indicates the median, and red asterisks mark the average values.

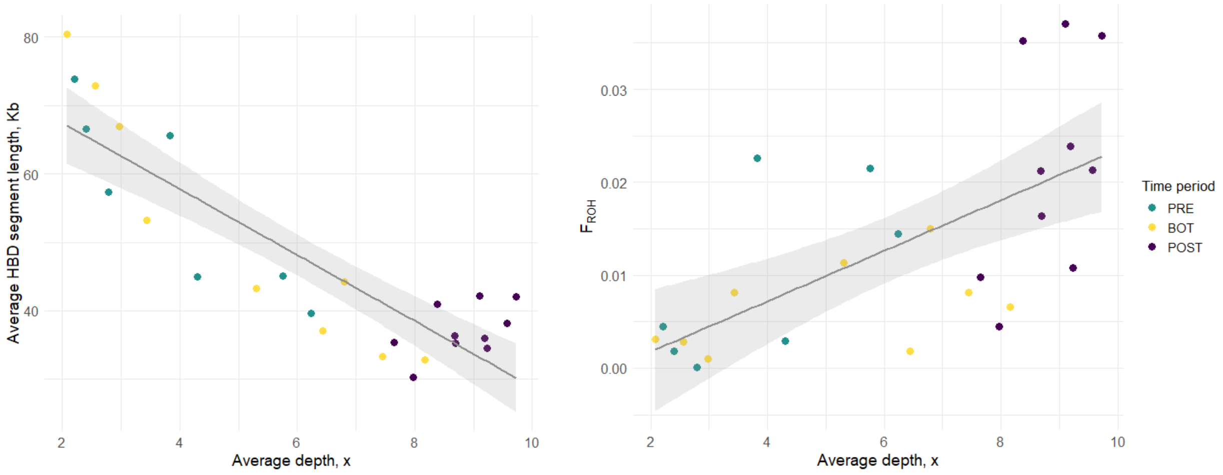


Figure S18. Correlation of runs of homozygosity (ROH) analysis parameters with average sequencing depth for Finnish golden eagles from three temporal groups: pre-bottleneck (PRE), bottleneck (BOT), and post-bottleneck (POST). The analysis is **based on 4,816,564 SNPs (full depth, including transitions). Left:** Average homozygosity-by-descent (HBD) segment length VS average sequencing depth (×). **Right:** Inbreeding coefficient estimated from ROH ($F_{ROH}$) VS average sequencing depth (×). Each dot is an individual, and colors correspond to temporal groups. The grey solid line is a linear trendline, and the shaded areas are 95% confidence intervals.

### DEMOGRAPHIC HISTORY

Demographic patterns inferred from the site frequency spectrum were generally consistent across datasets, although some differences were observed between downsampled and full-depth data (Table S3). In all datasets, PRE and BOT showed L-shaped SFS, whereas POST exhibited a relative loss of singletons. However, in the full-depth dataset, the POST SFS retained an L-shape (Figure S19). To validate the observed pattern of reduced number of singletons, we performed an independent downsampling replicate and re-analyzed the SFS metrics. Re-analysis resulted in consistent SFS shapes and fewer singletons in the POST group relative to other minor allele frequency classes.

Tajima's D differed significantly among temporal groups in all datasets (Table S3), but the direction of change varied. In the downsampled data, Tajima's D increased from PRE to POST, consistent with recent population contraction. In contrast, the full-depth dataset showed positive Tajima's D in all groups, with a slight decrease toward POST.

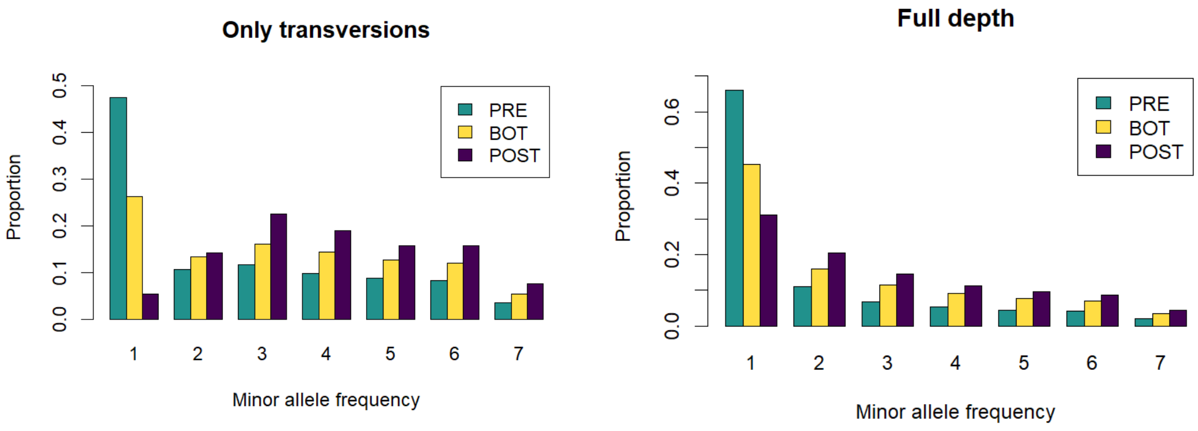


Figure S19. Folded site frequency spectrum (SFS) distributions of variable sites in Finnish golden eagles across three temporal groups: pre-bottleneck (PRE), bottleneck (BOT), and post-bottleneck (POST), **based on the downsampled dataset without transitions (left) and full-depth dataset with transitions (right)**. Each bar shows the number of sites with a given minor allele frequency class, i.e., the first column shows singletons (minor allele count = 1), followed by doubletons and higher-frequency classes up to alleles found in all seven individuals of a group. Note different Y-axis scales.
